# *Campylobacter jejuni* trigger signaling through host cell focal adhesions to inhibit cell motility and impede wound repair

**DOI:** 10.1101/2021.05.14.443831

**Authors:** Courtney M Klappenbach, Nicholas M. Negretti, Jesse Aaron, Teng-Leong Chew, Michael E. Konkel

**Affiliations:** School of Molecular Biosciences, College of Veterinary Medicine, Washington State University, Pullman, WA, USA, 99164-7520; Advanced Imaging Center, Janelia Research Campus, Howard Hughes Medical Institute, Ashburn, VA 20147

**Author notes:** Corresponding author: Michael E. Konkel, School of Molecular Biosciences, Biotechnology Life Sciences Building, Room 447, College of Veterinary Medicine, Washington State University, Pullman WA 99164.

**Keywords:** Pathogenesis, Focal adhesion modulation, Focal adhesion turnover, Cell migration, iPALM, bacteria-host cell interactions

## Abstract

*Campylobacter jejuni* is a major foodborne pathogen that exploits the focal adhesions of intestinal cells to promote invasion and cause severe gastritis. Focal adhesions are multiprotein complexes involved in bidirectional signaling between the actin cytoskeleton and the extracellular matrix. We investigated the dynamics of focal adhesion structure and function in *C. jejuni* infected cells using a comprehensive set of approaches, including confocal microscopy of live and fixed cells, immunoblots, and super-resolution iPALM. We found that *C. jejuni* infection of epithelial cells results in an increased focal adhesion size and altered topology. These changes resulted in a persistent modulatory effect on the host cell focal adhesion, as evidenced by an increase in cell adhesion strength, a decrease in individual cell motility, and a reduction of collective cell migration. We discovered that *C. jejuni* infection causes an increase in phosphorylation of paxillin and an alteration of paxillin turnover at the focal adhesion, which together represent a potential mechanistic basis for altered cell motility. Finally, we observed that infection of epithelial cells with the *C. jejuni* wild-type strain in the presence of protein synthesis inhibitor, a *C. jejuni* CadF and FlpA fibronectin-binding protein mutant, or a *C. jejuni* flagellar export mutant blunts paxillin phosphorylation and partially re-establishes individual host cell motility and collective cell migration. These findings provide a potential mechanism for the restricted intestinal repair observed in *C. jejuni*-infected animals and raise the possibility that bacteria targeting extracellular matrix components can alter cell behavior after binding and internalization by manipulating focal adhesions.

## INTRODUCTION

Significant research effort has focused on understanding how infectious agents cause disease in the host. Progress in understanding the fundamental mechanisms of bacterial pathogens has revealed common virulence strategies. Some of the most common targets for cellular level manipulation are the cytoskeleton and signaling pathways ^1; 2; 3^, which often enables bacterial pathogens to invade host cells ^4^. Promoting bacterial internalization can protect pathogens from the immune system and prolong infection ^5^. Many extensively studied pathogens, such as *Salmonella, Shigella, Vibrio*, and *Listeria*, have been found to use bacterial adhesins and secreted proteins (effectors) to alter signaling pathways, resulting in cytoskeleton rearrangements that promote their internalization ^1; 2; 5^. There are several major protein complexes connecting a cell to its external environment that can be targeted by bacteria for signaling manipulation.

In polarized epithelial cells, such as those in the intestine, there are four types of intracellular junctions: tight junctions, adherens junctions, hemidesmosomes, and focal adhesions. Tight and adherens junctions connect cells, whereas hemidesmosomes and focal adhesions facilitate cell-matrix connections ^6; 7^. Focal adhesions are unique from the others in that they connect the actin cytoskeleton to the extracellular matrix (ECM), making them dynamic and complex structures. The primary functions of focal adhesions are signaling, coordinating motility/migration, and adherence to the basal membrane. Signaling constitutes a major role for focal adhesions, as they can transmit bidirectional signals across the cell membrane ^8^. A vast range of information is transmitted through focal adhesion signaling, including signals for survival, proliferation, differentiation, platelet aggregation, embryo development, and sensing extracellular and intracellular tension ^7; 9; 10; 11^. A major function of these signals is to direct migration of epithelial cells in the intestinal villi, an important process for homeostasis ^12^. In the intestine, new cells are produced from pluripotent stem cells in the villus crypt, and these migrate along the villus-crypt axis towards the villus tip where they are eventually extruded ^12; 13^. The majority of studies on the cellular migration process have focused on collective epithelial cell migration in response to villi damage ^14; 15; 16; 17; 18^. When the villi become damaged, the basal lamina may become exposed or denuded ^14^. To repair this, healthy epithelial cells surrounding the wound will depolarize and migrate to cover the basal membrane ^14; 15; 16; 17; 18^. Cell migration is dependent on focal adhesion dynamics ^19^, whereby cells must simultaneously assemble new focal adhesions at the leading edge of the cell and disassemble old focal adhesions from the trailing edge of the cell ^20^. The balanced process of these actions allows cells to have directed migration ^20^. In addition to motility and migration, focal adhesions physically adhere cells to their external environment by attaching to the ECM. Focal adhesions depend on many proteins to manage these roles and functions.

Focal adhesions are composed of an organized array of proteins, collectively known as the focal adhesion “adhesome” ^21^. Integrins, transmembrane proteins that bind both ECM components and intracellular signal transduction proteins, comprise the base of focal adhesions ^8^. Structural proteins, such as talin and vinculin, connect integrins to the actin cytoskeleton and are required to strengthen focal adhesions ^22^. Last, signaling proteins allow the focal adhesion to send bidirectional signals through the cell membrane. These proteins include adaptors, scaffolds, kinases, and proteases ^10^. The coordinated action of adhesome proteins allows the cell to transmit signals between the extracellular environment and the intracellular actin cytoskeleton, as well as perform important cell functions such as motility/migration and adhesion.

Since many pathogens rely on the actin cytoskeleton and signaling proteins to cause infection in a host, focal adhesions are prime targets. *Campylobacter jejuni* is a major foodborne pathogen most commonly acquired from ingestion of undercooked chicken or foods cross-contaminated with raw poultry. Despite being the most common culture-proven cause of bacterial gastroenteritis worldwide ^23^, our knowledge of the specific virulence mechanisms and host-cell interactions is incomplete. Known is that severe *C. jejuni* infection results in diarrhea with blood in the stool and is accompanied by villus blunting ^24; 25; 26^. It is expected that villi restitution and collective cell migration would repair the tissue damage; however, it is not known if the phenotype observed in *C. jejuni* infection is due to slowing this process of villi repair. The purpose of this study was to determine if *C. jejuni* reduces cell motility, and if it does, the cellular mechanism that drives the alteration in cell behavior. We discovered that *C. jejuni* manipulates the dynamics of the focal adhesion, in part, via the CadF and FlpA fibronectin-binding proteins and secreted effector proteins. These findings constitute a new *C. jejuni*-driven cellular phenotype.

## RESULTS

### *C. jejuni* reduces host cell motility

To understand how *C. jejuni* manipulates the behavior of host cells during infection, we first examined the effect of *C. jejuni* on individual host cell motility using the A549 epithelial cell line. This cell line was chosen for the initial assays because the cells are highly motile ^27^. Cells were infected with *C. jejuni*, imaged for five hours, and then individual cells were tracked to quantify their motility. Cell motility was significantly decreased by infection with *C. jejuni* (Figure 1). Observation of the paths of individual cells over the course of the five hour infection shows that *C. jejuni* infected cells traveled a shorter distance (path distance) and remained closer to their starting position (path displacement) (Figure 1A-B). Processivity, path distance divided by path displacement, was calculated to quantify this observation. This measurement indicates a cell’s directional movement and is commonly used to assess cell motility ^28^. *C. jejuni* infected cells showed a significant decrease in processivity compared to non-infected cells (Figure 1C).

**Figure 1:**
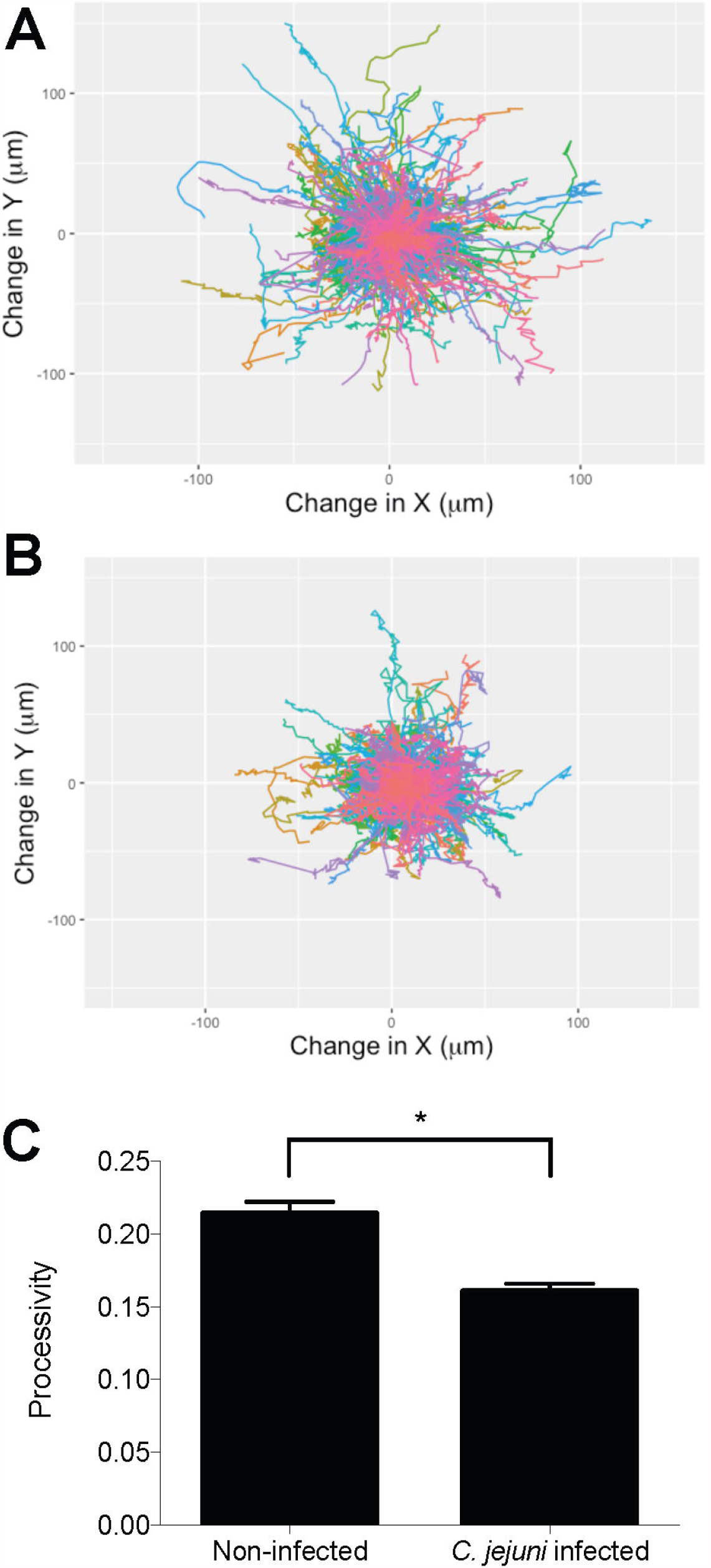
*C. jejuni* infection of A549 epithelial cells decreases their motility. A549 cells were grown on a lamina-based extracellular matrix prior to infection with *C. jejuni*. An image was taken every five minutes for five hours, and images were analyzed to track cell movement. Windrose plots (Panels A and B) show the tracked paths of individual cells (different colors) over five hours. **A**. Windrose plot of non-infected cells shows that most cells move far from the starting position (center of the plot: 0,0). **B**. Windrose plot of *C. jejuni* infected cells shows that they do not move as far from their starting position when compared to non-infected cells. **C**. *C. jejuni* causes a significant decrease in processivity, calculated as path distance divided by displacement averaged for all cells. Error bars represent SEM, * indicates *p* < 0.0001 (Students T Test). More than 500 cells were imaged per condition.

We also investigated the effect of *Salmonella* Typhimurium and *Staphylococcus aureus* on cell motility. *S. aureus* is a non-motile bacterium and causes sickness by staphylococcal enterotoxins (SE), triggering vomiting and inflammation of the intestine ^29^. *S*. Typhimurium, however, is a motile bacterium and causes disease by invading host cell intestinal tissues. Both *C. jejuni* and *Salmonella* infection require intimate bacterial-host cell contact for disease, while *S. aureus* does not. We investigated the effect of both *S*. Typhimurium and *S. aureus* on single cell motility. A549 epithelial cells were infected with the bacteria for one hour, and then the cells were rinsed and imaged for five hours. Chloramphenicol was used at a bacteriostatic concentration to ensure the bacterial CFU was consistent throughout the experiment. Cells infected with *S*. Typhimurium had significantly lower motility than non-infected cells; however, there was no significant difference in cell motility for cells infected with *S. aureus* (Supplemental Figure 1). Our findings are consistent with the fact that *S*. Typhimurium binds to fibronectin and induces the formation of focal adhesion-like complexes during cell invasion ^30; 31^. This result suggests that the inhibition of host cell motility is conserved among some bacterial species.

### *C. jejuni* increases the size of the focal adhesion in a flagellar dependent manner

Since epithelial cell motility is driven by a balance between focal adhesion assembly and disassembly ^20^, we investigated if *C. jejuni* alters host cell focal adhesion structure. The human INT 407 cell line was used for these assays, as this particular cell line has been used extensively by researchers to dissect *C. jejuni*-host cell interactions ^32; 33; 34^. Cells were infected with *C. jejuni* and then fixed and stained for the host cell protein paxillin, which is a major signaling and adaptor protein in focal adhesions ^35^. Confocal microscopy showed significant changes in paxillin localization between infected and non-infected cells (Figure 2A). In non-infected cells, paxillin was largely localized in the cytosol, whereas in *C. jejuni* infected cells it was concentrated at focal adhesions. This change was found to be significantly different when scored by an individual blinded to the sample identity, where a score of 0 represented complete cytosolic localization and 2 represented no cytosolic localization. The individual determined the score for each cell in the randomized, blinded images using a scoring key. Consistent with this observation, quantitation of focal adhesion size in infected and non-infected cells showed that *C. jejuni* caused a significant increase in focal adhesion size from 30 to 75 minutes post-infection (Figure 2B). Many *C. jejuni* also appeared to localize near focal adhesions (Figure 2D-E). This is consistent with previously published results ^36^ showing that *C. jejuni* colocalized with paxillin and vinculin, another major focal adhesion protein. This led us to investigate the bacterial factors required for the observed changes.

**Figure 2:**
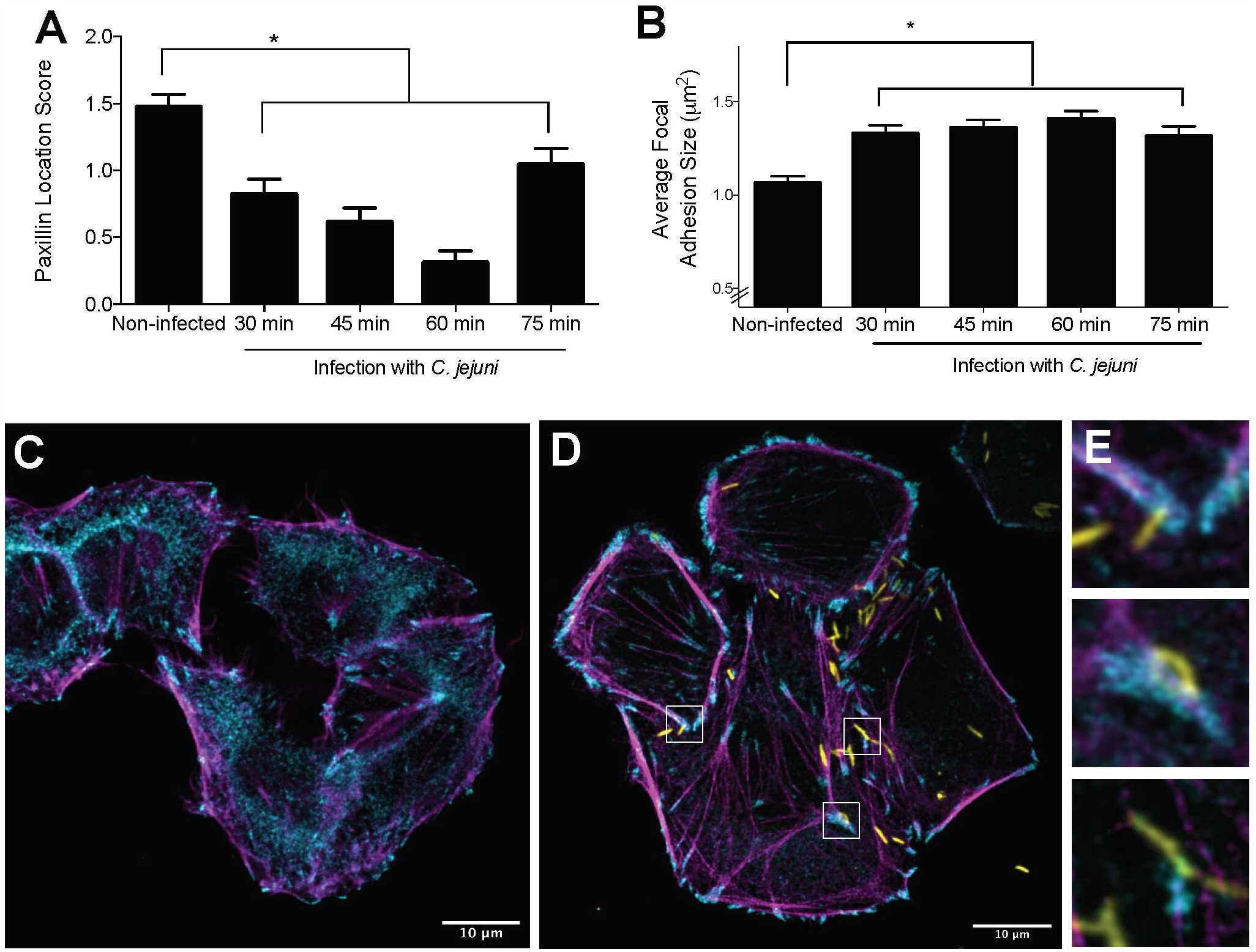
*C. jejuni* associates with and increases the size of the focal adhesion footprint and changes the localization of paxillin. INT 407 cells were infected with *C. jejuni* for 30 to 75 minutes prior to fixing and immunostaining for paxillin, actin, and *C. jejuni*. **A**. Paxillin is localized to the cytosol in non-infected cells and localized at the focal adhesion in *C. jejuni* infected cells. Paxillin localization was determined by an individual blinded to the samples, where 0 = complete focal adhesion localization and 2 = cytosolic localization. Error bars represent SEM of > 46 cells, and the * indicates *p* < 0.05 compared to the non-infected sample (One-way ANOVA, Dunnett’s multiple comparisons test). **B**. *C. jejuni* causes a significant increase in focal adhesion size at 30, 45, 60, and 75 minutes post-infection. Error bars represent standard deviation of > 1000 focal adhesions, and the * indicates *p* < 0.05 compared to the non-infected sample (One-way ANOVA, Tukey’s multiple comparisons test). **C**. Representative image of non-infected cells. **D**. Representative images of *C. jejuni* infected cells. **E**. Insets from panel D showing that *C. jejuni* is associated with paxillin. Paxillin is shown in cyan, actin in magenta, and *C. jejuni* in yellow. The focal adhesions in infected cells appear larger than those in non-infected cells, and there is less paxillin in the cytosol of *C. jejuni* infected cells.

The flagellum is an important virulence factor for *C. jejuni*; it confers motility and also serves as the bacterium’s type three secretion system to deliver effector proteins into a host epithelial cell ^37; 38; 39; 40; 41^. To determine if the changes in focal adhesion structure were driven by bacterial factors, a *C. jejuni* Δ*flgL* mutant was tested alongside wild-type bacteria. The *flgL* gene encodes the flagella hook junction protein, and a *C. jejuni* Δ*flgL* mutant is non-motile and incapable of protein export including secretion of the effectors ^32; 42^. In contrast to the *C. jejuni* wild-type strain, the Δ*flgL* mutant had no effect on focal adhesion size, showing a similar focal adhesion distribution to non-infected cells (Figure 3). The *C. jejuni flgL* complemented isolate, however, restored the phenotype observed in the wild-type bacteria. This result demonstrates that a functional flagellum (*i.e*., secretion apparatus) is required to increase the size of the focal adhesion.

**Figure 3:**
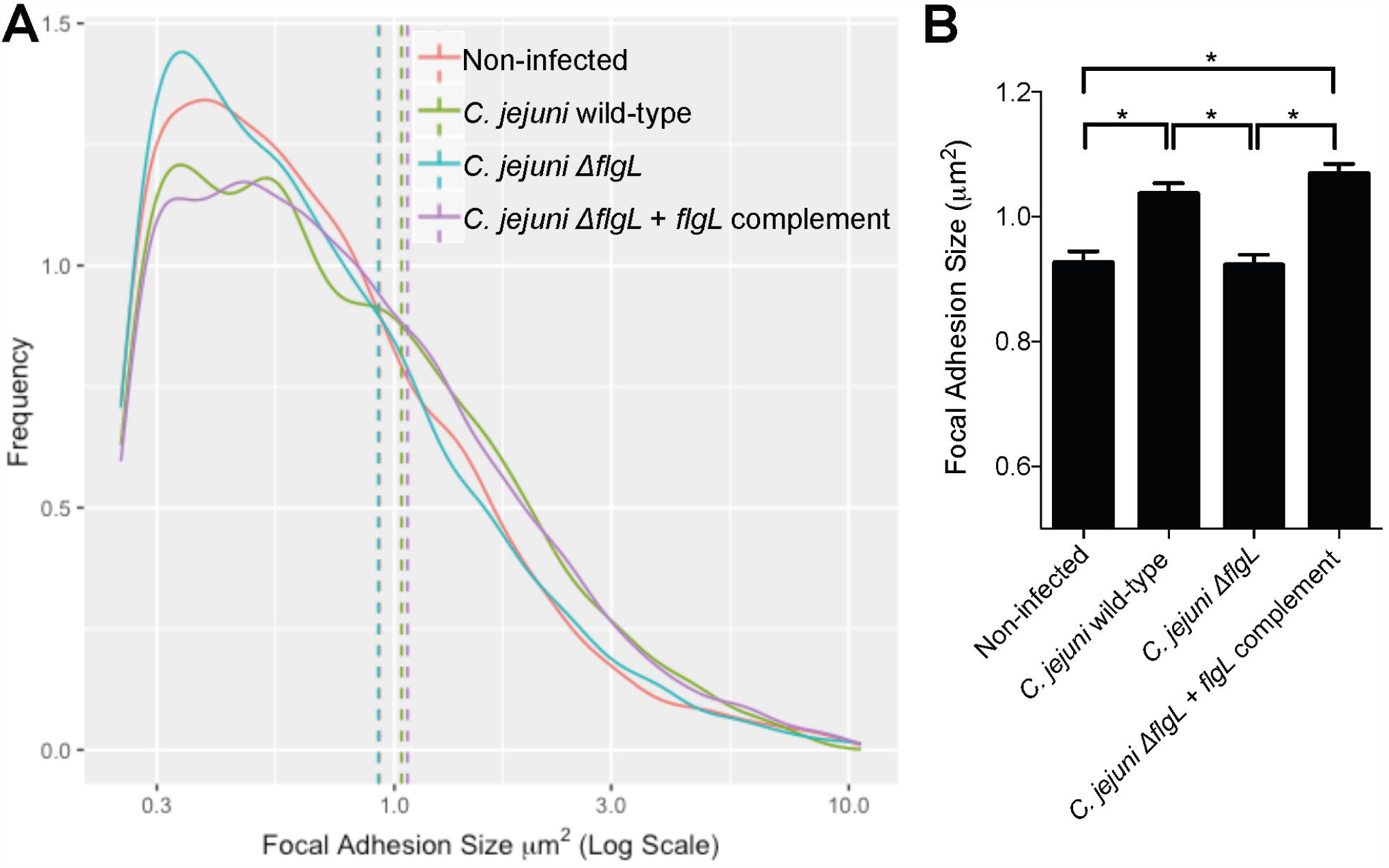
*C. jejuni* driven focal adhesion size increase is dependent on a functional flagellum. INT 407 cells were infected with a *C. jejuni* wild-type strain, a *C. jejuni* Δ*flgL* mutant, or a *C. jejuni* Δ*flgL + flgL* complemented strain for 60 minutes prior to fixing and immunostaining for paxillin, actin, and *C. jejuni*. Focal adhesion size was determined by measuring the paxillin footprint. **A**. Frequency distribution of focal adhesion sizes for the four conditions is shown. Solid lines show frequency distribution, and dashed lines show the mean of all focal adhesions within a category. Non-infected and *C. jejuni* Δ*flgL* infected cells have a similar distribution of focal adhesion sizes while cells infected with the *C. jejuni* wild-type strain and *C. jejuni* Δ*flgL + flgL* complemented isolate have a similar distribution. **B**. The bar graph shows the mean focal adhesion size for each condition. The *C. jejuni* wild-type strain increases the average focal adhesion size when the *flgL* gene is present, indicating that motility and/or protein secretion are required. Error bars represent SEM, * indicates *p* < 0.0001 (One-way ANOVA, Tukey’s multiple comparisons test, > 3000 focal adhesions measured per condition).

To further explore how *C. jejuni* manipulates the focal adhesion, we tested single cell motility using a Δ*cheB* mutant and Δ*flhF* mutant that had been generated previously ^32^. The *C. jejuni* Δ*flgL* mutant was included as a control. The *C. jejuni* CheB protein has methylesterase activity and CheR has methyltransferase activity. Both are involved in chemotaxis ^43^. Deletion of the *cheB* gene has a polar effect on *cheR* expression, therefore, we will refer to this isolate as a *cheBR* mutant. Notable is that the *cheBR* mutant has a functional flagellum (*i.e*., secretion apparatus, as evident by the presence of the flagellum), is motile, and does not exhibit a defect in adherence or invasion when compared to a wild-type strain ^32^. The *C. jejuni* FlhF protein binds to the promoters of flagellar genes and regulates their transcription. Disruption of the *flhF* gene in *C. jejuni* alters the expression of flagellar genes and the process of flagellar biosynthesis, resulting in a non-flagellar phenotype, absence of motility, and decreased epithelial cell adherence and invasion ^32^. The *cheBR* mutant was able to limit A549 host cell motility to a level similar to infection of cells with the wild-type bacteria (Supplemental Figure 2). In contrast, A549 cells infected with the Δ*flhF* mutant or Δ*flgL* mutant were more motile than cells infected with the wild-type isolate (Supplemental Figure 2). Although a subset of flagellar proteins needed for motility are necessary for a functional export apparatus ^42^, these data suggest that bacterial motility and/or a functional flagellar export apparatus is required to alter (reduce) host cell motility.

### *C. jejuni* slows paxillin turnover and changes the topology of the focal adhesion

To determine the mechanism of the focal adhesion size increase, we investigated the turnover rate of paxillin at the focal adhesion in INT 407 cells. Proteins at the focal adhesion regularly turnover and have unique residency times ^21; 44^. INT 407 cells were transfected with a plasmid containing tdEos conjugated to paxillin. The tdEos protein photoswitches from green to red when exposed to UV light. By photoswitching one-half of a cell’s focal adhesions, it is possible to measure focal adhesion turnover by observing the rate that the photoswitched protein replaces the non-photoswitched protein over time (Figure 4A). At the non-photoswitched half of the cell, viewing the red channel only shows that red photoswitched paxillin increased over time at focal adhesions; this observation demonstrates that the photoswitched paxillin (red) migrates to the site of the non-photoswitched paxillin (green) (Figure 4B). Moreover, there was a significant decrease in the average rate of red fluorescence appearing in the non-photoswitched half of the cell for the *C. jejuni* infected cells compared to non-infected cells. This implies slower focal adhesion turnover and a decrease in the dynamicity of the focal adhesions. As a result of *C. jejuni* manipulating focal adhesion structure and signaling, the focal adhesion’s ability to turnover paxillin is slowed.

**Figure 4:**
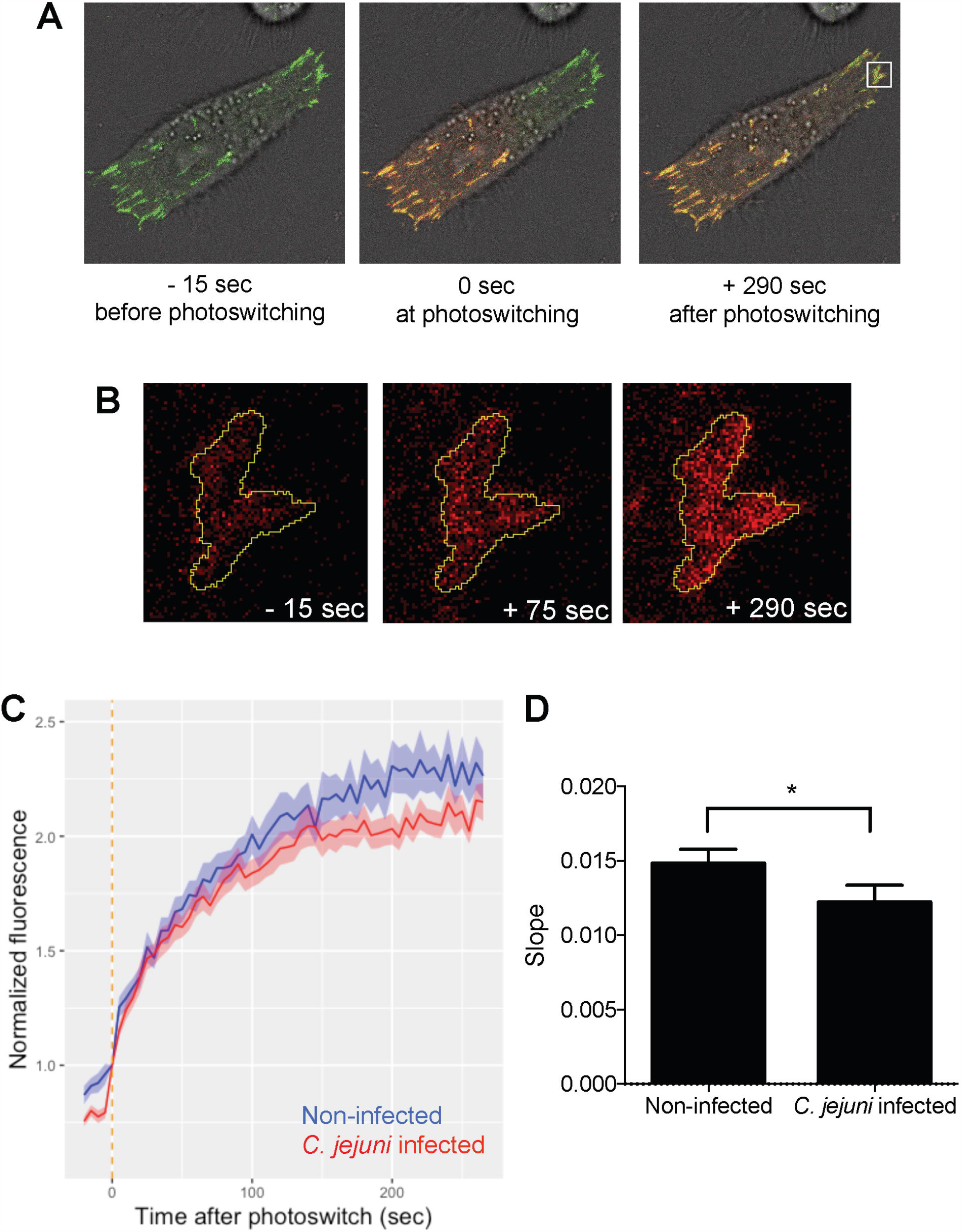
*C. jejuni* infection slows the turnover of paxillin at the focal adhesion. INT 407 cells were transfected with a tdEos-Paxillin plasmid, which photoswitches from green to red when exposed to UV light. Cells were infected for approximately 60 minutes with *C. jejuni* prior to imaging. **A**. One-half of the cell was exposed to UV light to switch the tdEos-paxillin from green to red, and then the cell was imaged every 5 seconds for approximately 300 seconds. As the focal adhesion turns over, the green paxillin in the non-photoswitched half of the cell is replaced with red paxillin from the photoswitched half of the cell. **B**. Enlarged view of a focal adhesion in the non-photoswitched half of the cell showing the red fluorescence channel only increasing over time. **C**. Normalized fluorescence was calculated as the average intensity of red fluorescence at focal adhesions in the non-photoswitched half of the cell divided by each focal adhesion’s initial red fluorescence intensity. Lines represent the average of all focal adhesions for a condition. The focal adhesions of *C. jejuni* infected cells took longer to show increased red fluorescence and reached an overall lower red fluorescence intensity. The vertical orange line represents the time of photoswitching. **D**. The average slope of red fluorescence in the non-photoswitched half of the cell was calculated for all focal adhesions by linear regression between normalized fluorescence and time. *C. jejuni* infected cells had decreased slope compared to non-infected cells, indicating a slower rate of paxillin turnover. Error bars represent SEM and * indicates *p* < 0.0001 (Student’s T test). More than 200 focal adhesions measured per condition.

While we observed that *C. jejuni* infection of epithelial cells caused an increase in focal adhesion size, it was not clear if the structural topology within the focal adhesion was changing. To observe the micro-scale structure of the focal adhesions in *C. jejuni* infected cells, super-resolution iPALM (interferometric photoactivated localization microscopy) was utilized to observe the organization of proteins within the focal adhesion at sub-20 nm resolution ^45^. tdEos-Paxillin was used as our indicator of the focal adhesion structure due to its critical role in focal adhesion organization and *C. jejuni* invasion. We observed that the spatial localization of paxillin was significantly different in *C. jejuni* infected cells compared to non-infected cells (Figure 5, Supplemental Figure 3). The average Z position of paxillin, measured as the distance from the gold fiducial beads embedded in the coverslip, was 40.3 nm ± 2.2 nm in non-infected cells. This is consistent with the published value of 36.0 nm □±□ 4.7□nm ^46^. Paxillin in *C. jejuni* infected cells was significantly higher, at 56.7 nm ± 1.6 nm (Figure 5A).□ In addition, we calculated the thickness of the paxillin plaque. The plaque in non-infected cells had an average thickness of 26.9 nm ± 1.8 nm, while *C. jejuni* infected cells had significantly thicker focal adhesions of 32.6 nm ± 0.8 nm (Figure 5B). These results supported the confocal microscopy studies and demonstrated that *C. jejuni* manipulates the nanoscale localization of paxillin in the focal adhesion.

**Figure 5:**
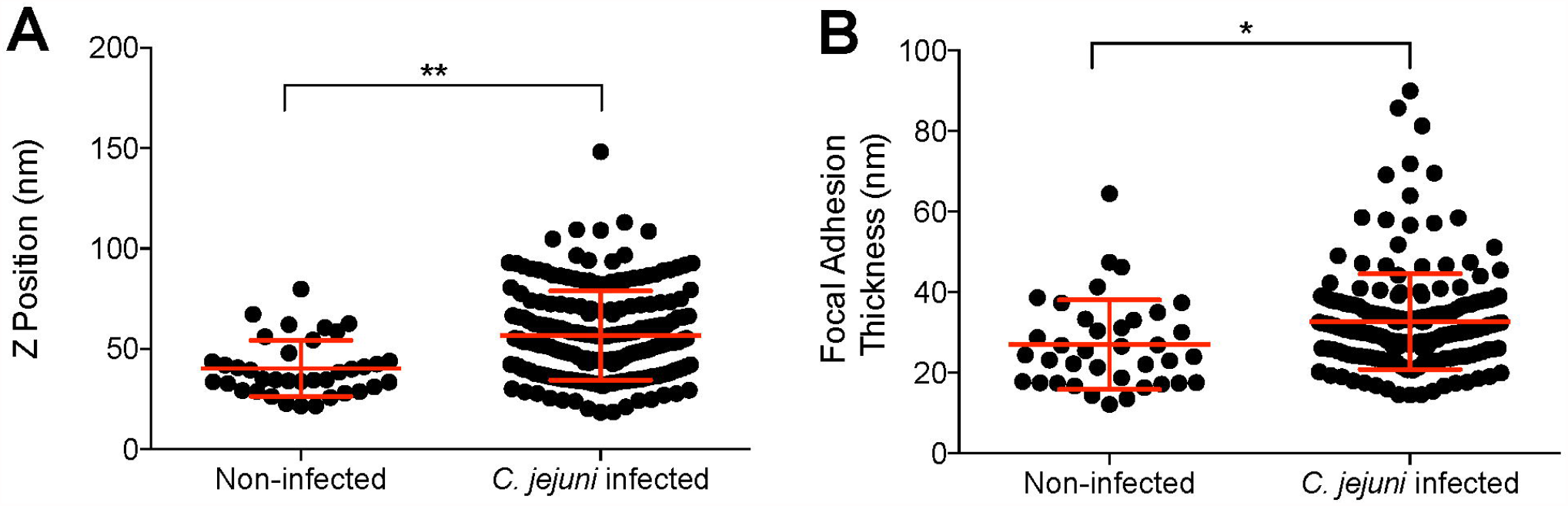
*C. jejuni* changes the nanoscale topology of the focal adhesion plaque. Super-resolution iPALM microscopy was performed to determine the nanoscale architecture of the focal adhesion in *C. jejuni* infected cells compared to non-infected cells. **A**. The Z position of a paxillin plaque was determined as the average z position of all fluorophores detected within a focal adhesion. The average Z position of *C. jejuni* infected focal adhesions was significantly higher than non-infected cells. **B**. The average thickness of the focal adhesion was determined as the inner quartile range of all fluorophores within the focal adhesion. The average thickness of *C. jejuni* infected focal adhesions was significantly higher than non-infected cells. Error bars represent SD, * indicates *p* < 0.01, ** indicates *p* < 0.0001 (Student’s T-Test).

### *C. jejuni* enhances signaling pathways at the focal adhesion

Phosphorylation of paxillin (at Tyr118) is indicative of focal adhesion assembly and activation of the FAK and Src kinases ^47; 48^. Previous research has revealed that paxillin is phosphorylated during *C. jejuni* infection of INT 407 cells in a time-dependent manner ^49^. To determine if paxillin phosphorylation is related to focal adhesion signaling, we tested the role of focal adhesion kinase (FAK) and Src kinase in paxillin phosphorylation and *C. jejuni* invasion. It is well established that these two kinases form a complex together at focal adhesions to phosphorylate many proteins, including paxillin ^50^. We treated INT 407 epithelial cells with selective FAK (TAE226) and Src (PP2) inhibitors prior to and during infection with *C. jejuni*. Paxillin was then immunoprecipitated from the cells, and the relative amount of phosphorylated paxillin was determined by immunoblot analysis. Consistent with previous research ^49^, *C. jejuni* caused a significant increase in phosphorylated paxillin. Importantly, the presence of the TAE226 or PP2 inhibitors eliminated this effect (Figure 6A and 6B). To verify the biological significance of these kinases, *C. jejuni* invasion in the presence of these inhibitors was determined by the gentamicin protection assay. Both kinase inhibitors caused a significant reduction in the number of internalized bacteria (Figure 6C), which is consistent with previous reports ^36; 51; 52; 53^. To further confirm that the changes in paxillin signaling are related to focal adhesion dynamics, we investigated the phosphorylation of paxillin at the focal adhesion by microscopy. INT 407 epithelial cells were infected with *C. jejuni*, then fixed and stained with paxillin and phosphorylated paxillin (Y118) antibodies. Cells were imaged by confocal microscopy, and the intensity of paxillin and phosphorylated paxillin was quantified at each focal adhesion. The relative amount of phosphorylated paxillin compared to total paxillin increased significantly in *C. jejuni* infected cells compared to non-infected cells (Figure 6D-G). This result demonstrated that the *C. jejuni-*driven paxillin signaling changes are occurring at the focal adhesion.

**Figure 6:**
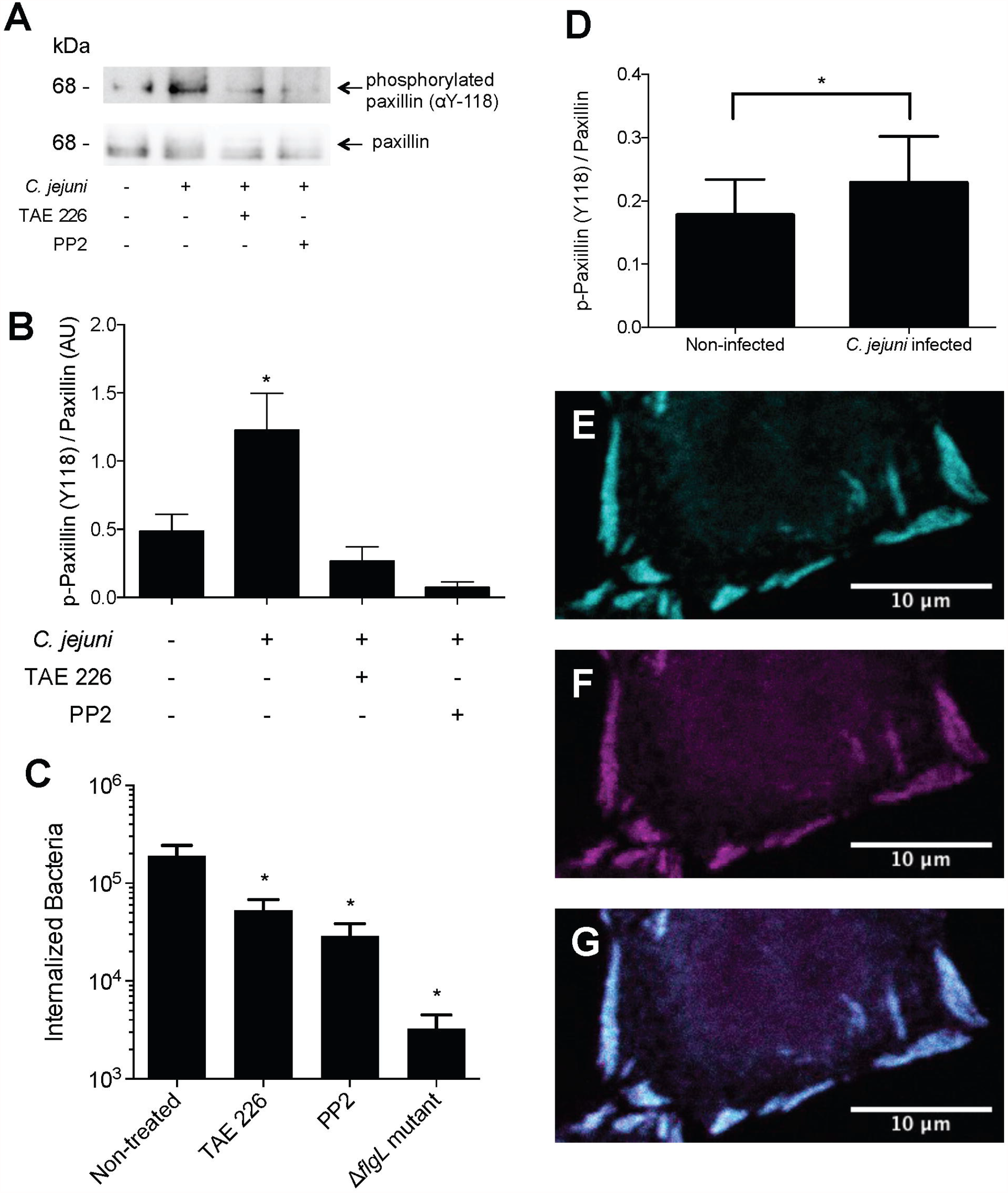
*C. jejuni* invasion results and paxillin phosphorylation by FAK and Src kinases. **A**. INT 407 cells were incubated with *C. jejuni* for 45 minutes in the presence of TAE226 (an inhibitor of FAK) or PP2 (an inhibitor of Src). Cells were lysed, and paxillin was immunoprecipitated. SDS-polyacrylamide gels were run, and blots were probed for phosphorylated (Y118) paxillin and total paxillin. **B**. Band intensity was measured, and phosphorylated paxillin was normalized by total paxillin. Results showed that *C. jejuni* causes a significant increase in paxillin phosphorylation, but that the addition of TAE226 or PP2 eliminates this effect. Error bars represent standard deviation between two biological replicates. **C**. INT 407 cells were incubated with *C. jejuni* and inhibitors prior to determining the CFU of internalized bacteria by the gentamicin protection assay. TAE226 and PP2 caused a significant decrease in the number of internalized bacteria compared to non-treated cells. Error bars represent standard deviation between technical replicates. * indicates *p <* 0.05 compared to non-treated or non-infected wells (One-way ANOVA, Dunnett’s multiple comparisons test). Δ*flgL* mutant (non-invasive) was used as a negative control. **D-G**. INT 407 cells infected with *C. jejuni* harboring a GFP plasmid for 60 minutes show that phosphorylated paxillin is localized to the focal adhesion. Cells were then fixed and stained with a paxillin and phospho-paxillin (Y118) antibody. Focal adhesions were imaged by confocal microscopy. The relative amount of phospho-paxillin was determined by segmenting the focal adhesion and measuring the intensity of paxillin and phospho-paxillin within the segmented area. The phospho-paxillin intensity was divided by the paxillin intensity for each focal adhesion. **D**. The relative amount of phospho-paxillin at the focal adhesion increased significantly in *C. jejuni* infected cells compared to non-infected cells. Error bars represent the standard deviation of all focal adhesions (> 6000 total sites examined per condition) and * represents *p* < 0.0001 (Student’s T Test). **E**. Representative image of focal adhesions showing paxillin staining only. **F**. Focal adhesion showing phospho-paxillin staining only. **G**. Composite of E and F, showing paxillin and phospho-paxillin staining.

### Biological significance of focal adhesion manipulation

Focal adhesion size has been found to correlate with cellular adherence strength ^54; 55^. To see if *C. jejuni* is manipulating focal adhesion strength in addition to size, we used a trypsin-based cell detachment assay. Trypsin is a protease that causes cultured cell detachment. At low concentrations, trypsin treatment can be used to determine adhesion strength ^54^. Cells that are more strongly attached to the substrate will take longer to detach than cells more weakly attached. INT 407 cells were infected with *C. jejuni* or treated with nocodazole prior to treatment with a low concentration of trypsin. Nocodazole inhibits microtubule polymerization and was used as a positive control, as it increases focal adhesion size and strength ^56^. Cells were imaged for 15 minutes with a phase-contrast microscope to observe the cell rounding that precedes detachment. *C. jejuni* and nocodazole both caused a delay in cell rounding when compared to non-infected cells (Figure 7), indicating that they both caused an increase in focal adhesion strength.

**Figure 7:**
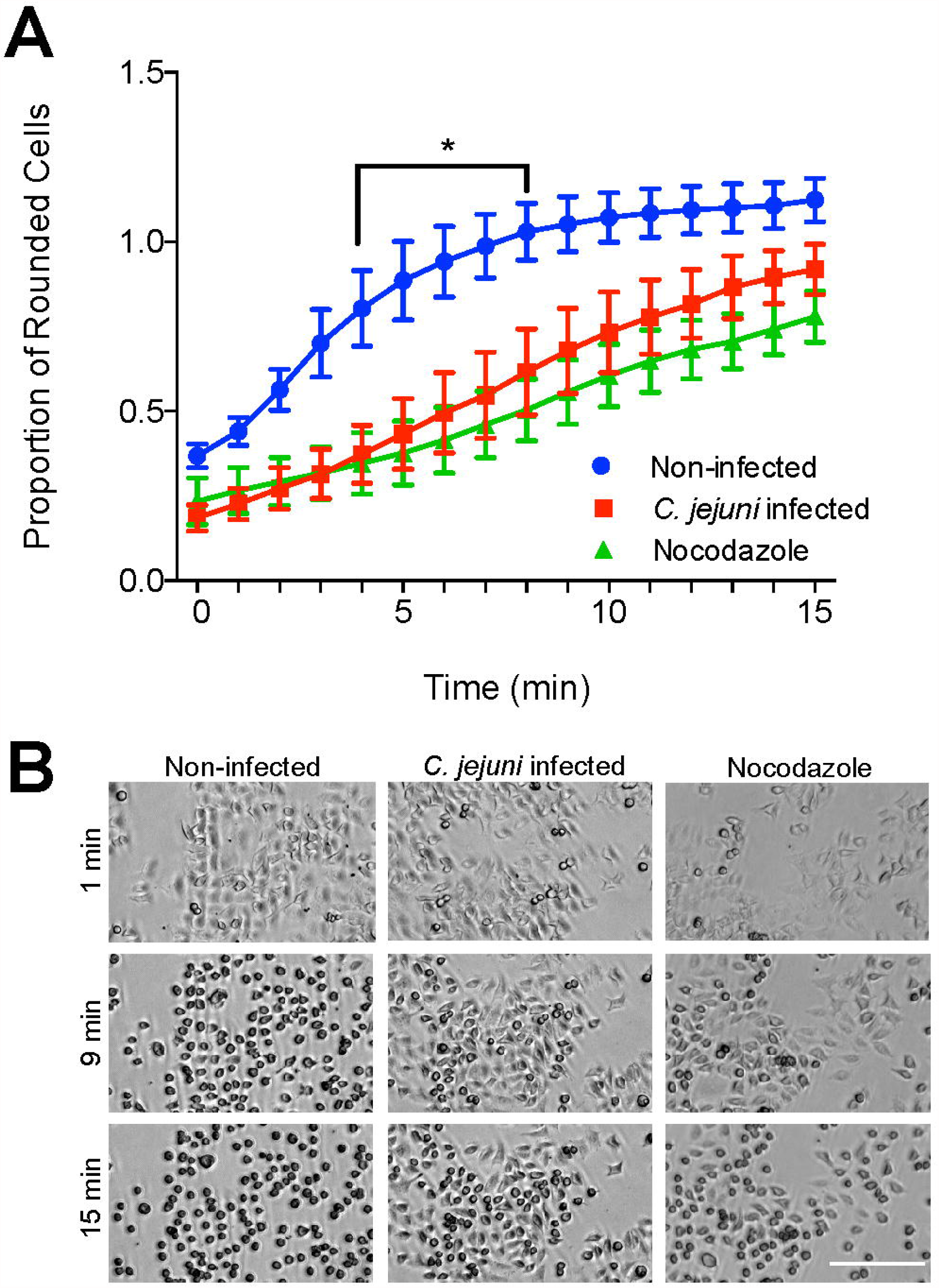
*C. jejuni* increases host cell adhesion strength. INT 407 cells were seeded 24 hours before infection with *C. jejuni* or treatment with 20 μM nocodazole (a microtubule polymerization inhibitor that increases focal adhesion size and strength) for 45 minutes. Cells were treated with a low concentration of trypsin to induce cell rounding and then imaged every 5 to 30 seconds for 15 minutes. **A**. Rounded cells were counted at each minute after trypsin treatment and divided by the total number of cells. The number of rounded cells appearing per minute was faster in non-infected cells than *C. jejuni* and nocodazole treated cells, indicating increased adhesion strength in these conditions. Error bars represent SEM, * indicated *p <* 0.05 between *C. jejuni* and non-infected conditions (Two-way ANOVA, Sidak’s multiple comparisons test). **B**. Images of cells treated with *C. jejuni*, nocodazole, or non-infected. In non-infected cells, most cells appear rounded at 9 minutes post-trypsin treatment, and all cells are rounded at 15 minutes. In nocodazole and *C. jejuni* infected conditions, fewer cells are rounded at 9 and 15 minutes. Scale bar represents 200 μm.

To understand the effect of focal adhesion manipulation on wound healing, we tested the collective cell migration of human T84 epithelial cells in response to *C. jejuni*. The human T84 colonic cell line represents an ideal model to mimic tissue-level responses to *C. jejuni*, as *C. jejuni* infects the human colon ^57; 58^. Additionally relevant to these experiments is that T84 cells have been used to mimic the wound healing response in the intestinal villi ^15^. Small circular scratches were made in an epithelial cell monolayer and imaged for 4 days to observe wound closure. There were no significant differences in cell number or percent viability between infected and non-infected conditions 6 days after scratching cells (data not shown). Non-infected monolayers healed almost completely in 4 days, while monolayers infected with *C. jejuni* did not (Figure 8). Linear regression analysis revealed that non*-*infected scratches healed at a rate of 0.0082 ± 0.00063 (normalized area/hour) while *C. jejuni* infected scratches healed significantly slower at 0.0042 ± 0.00046 (*p* < 0.0001, ANCOVA). The average initial wound size was not significantly different between samples (*p* = 0.315, Student’s T Test). This result shows that *C. jejuni* significantly decreases the ability of the epithelial cell monolayer to heal wounds.

**Figure 8:**
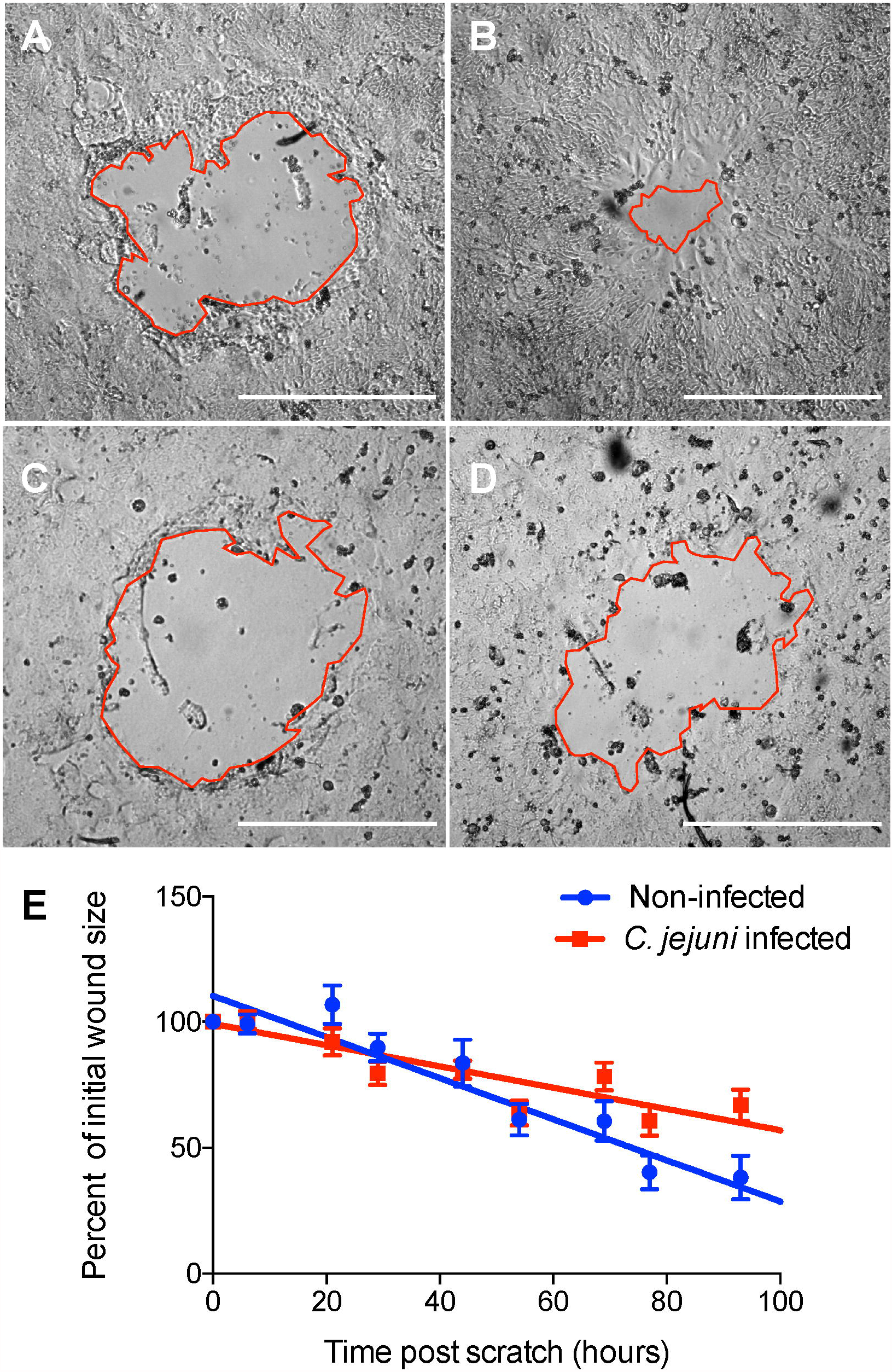
*C. jejuni* decreases the rate of epithelial wound healing. T84 cells were grown until confluency then scratched by aspiration to create a small circular wound. Cells were infected with *C. jejuni* for three hours and then imaged twice daily for 4 days to monitor wound closure. **A**. Non-infected scratch immediately after wounding. **B**. Non-infected scratch 100 hours after wounding. **C**. *C. jejuni* infected scratch immediately after wounding. **D**. *C. jejuni* infected scratch 100 hours after wounding. Red lines outline the wound, and bars represent 0.5 mm. **E**. Average percent of initial wound size over time. Lines represent linear regression of points for each condition. The rate of closure is significantly lower for *C. jejuni* infected scratches compared to non-infected scratches (Non-infected = - 0.0082 ± 0.00063, *C. jejuni* infected = - 0.0042 ± 0.00046, *p* < 0.0001, ANCOVA).

### *Mechanism of* C. jejuni *focal adhesion manipulation*

We have shown that *C. jejuni* manipulates the size, structure, and composition of focal adhesions. Based on previous research connecting *C. jejuni* adhesins to focal adhesion components ^49; 53; 59^, we hypothesized that the CadF and FlpA adhesins could be responsible for driving the initial changes in the focal adhesion and secreted effector proteins could contribute to the persistent modulatory effect of the focal adhesion. First, we tested if the CadF and FlpA adhesins were required for signaling changes in the focal adhesion by examining paxillin phosphorylation. INT 407 cells were infected with *C. jejuni* wild-type strain, *C. jejuni* lacking the *cadF* and *flpA* genes (Δ*cadF* Δ*flpA*), or the *C. jejuni* Δ*cadF* Δ*flpA* with the *cadF* and *flpA* genes restored (Δ*cadF* Δ*flpA + cadF flpA*). Cells were also infected with the *C. jejuni* Δ*flgL* mutant, as this isolate is deficient in the secretion of the Cia effector proteins. More specifically, the CiaD effector protein, which is not secreted from a flagellar mutant bacterium, activates the MAP kinase signaling pathway, including Erk 1/2, which is in a complex with paxillin ^60; 61^. Non-infected cells were used as a negative control. We observed a reduced level of phosphorylation in cells infected with the *C. jejuni* Δ*cadF* Δ*flpA* mutant when compared to cells infected with the *C. jejuni* wild-type strain (Figure 9A, Supplemental Figure 4). In addition, we observed a reduced level of phosphorylation in cells infected with the *C. jejuni* Δ*flgL* mutant when compared to *C. jejuni* wild-type strain (Figure 9A, Supplemental Figure 4). These findings suggest that the bacterial adhesins and secreted proteins work cooperatively to alter the paxillin activation and downstream signaling pathways.

**Figure 9:**
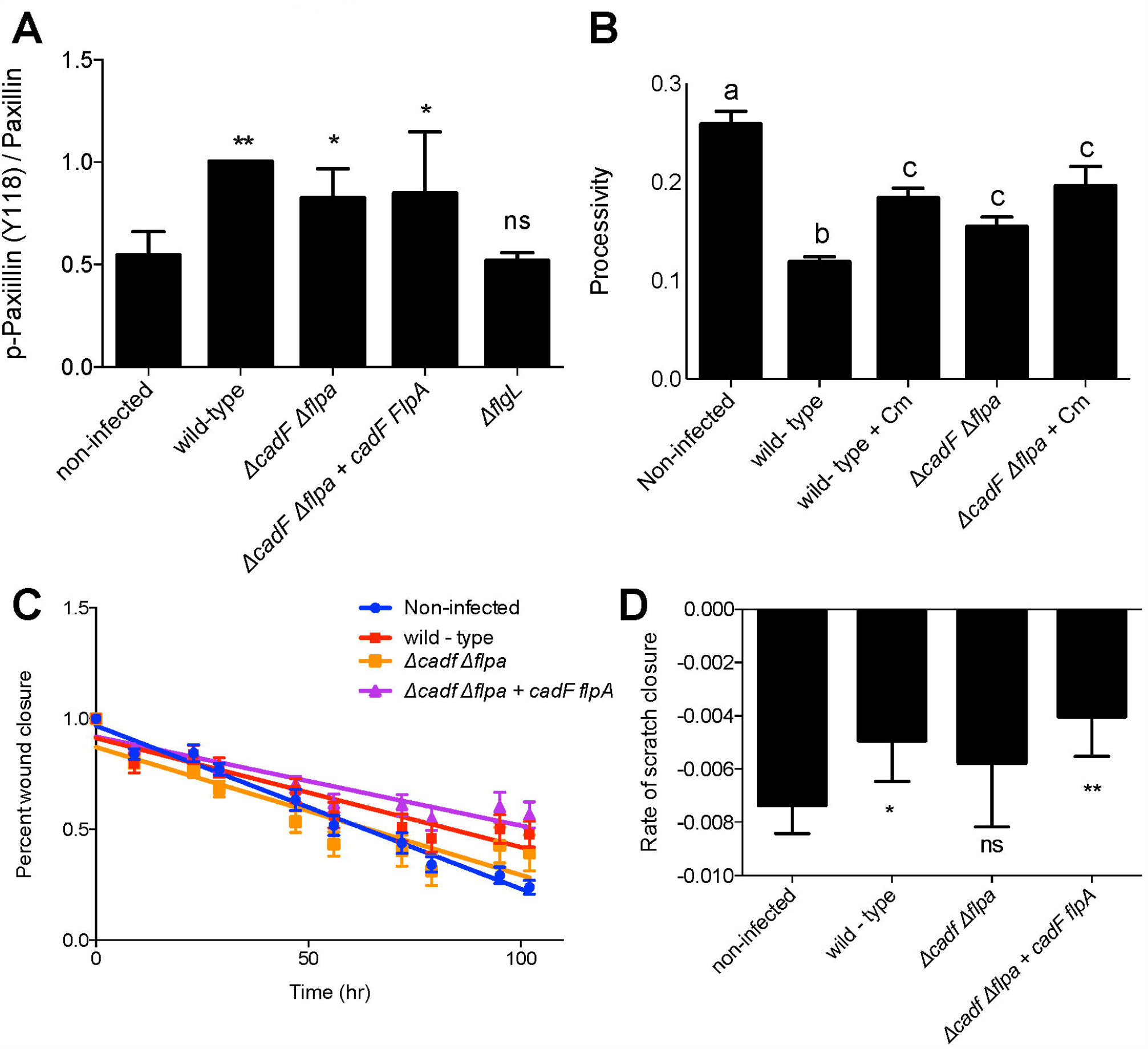
The *C. jejuni* CadF and FlpA adhesins contribute to focal adhesion modification. **A**. INT 407 cells were infected with *C. jejuni* for 60 minutes then lysed. Protein gels were run with whole-cell lysates and immunoblots were probed for total paxillin and phosphorylated paxillin (Y118). Densitrometry was performed to quantify band intensity and phosphorylated paxillin was normalized to total paxillin. 5 biological replicates were normalized to their wild-type condition and averaged. *C. jejuni* wild-type caused a significant increase in paxillin phosphorylation compared to the non-infected condition. The *C. jejuni* Δ*flgL* (flagella hook-junction protein, mutant is non-motile and protein secretion negative) demonstrated low levels of phosphorylation, and the Δ*cadF* Δ*flpA* mutant and complement had a modest reduction in phosphorylation compared to cells infected with the *C. jejuni* wild-type strain. Error bars represent SD, * indicates *p* < 0.5 and ** indicates *p* < 0.001 by one-way ANOVA with Dunnett’s multiple comparison test. **B**. Migratory A549 cells were grown on a laminin-based extracellular matrix overnight then infected with *C. jejuni* wild-type strain or *C. jejuni* Δ*cadF* Δ*flpA* mutant with and without chloramphenicol to inhibit protein synthesis (and the secretion of the bacterial effector proteins). Cells were imaged for 5 hours to observe migration over time. Processivity was calculated to represent directional migration. Wild-type bacterial had significantly decreased processivity, however migration was partially restored by the addition of chloramphenicol and when the *C. jejuni* Δ*cadF* Δ*flpA* mutant was used. Use of both the adhesion mutant and chloramphenicol resulted in the highest processivity of all infected conditions. Bars represent the average of > 40 cells, error bars represent SEM. By one-way ANOVA with Tukey’s multiple comparison’s test: # indicates significant difference from all other conditions (*p* < 0.05), † indicates significant difference from all other conditions (*p* < 0.05), ‡ indicates significant difference from non-infected and wild-type conditions (*p* < 0.05). **C**. T84 cells were grown to confluency then scratched by aspiration to create a small circular wound. Cells were left non-infected or infected with *C. jejuni* wild-type strain, *C. jejuni* Δ*cadF* Δ*flpA* deletion mutant, or the *C. jejuni* Δ*cadF* Δ*flpA* + *cadF flpA* complemented isolate. Scratched cells were imaged twice daily for four days. Non-infected and *C. jejuni* Δ*cadF* Δ*flpA* infected scratches demonstrated a faster closure of the scratch than *C. jejuni* wild-type and *C. jejuni* Δ*cadF* Δ*flpA* + *cadF flpA* infected cells. 7 – 12 scratches per condition, error bars represent SEM, one of 2 biological replicates. **D**. Quantification of the rate of scratch closure for *C. jejuni* infected cells. Rate was determined by finding the slope of the line describing normalized scratch area over time for each scratch and averaged. Error bars represent SD, and * indicates *p* < 0.01, and ** indicates *p* < 0.001 compared to the non-infected condition by one-way ANOVA with Dunnett’s multiple comparison test.

To determine how CadF and FlpA may contribute to functional changes to the focal adhesion, we examined their role in individual and collective cell migration. Individual cell migration was tested using A549 epithelial cells. Cells were grown on a laminin-based extracellular matrix, infected with the *C. jejuni* isolates, and imaged for 5 hours to observe migration. In addition to testing the dependence on the adhesins, we wanted to test if metabolically active bacteria were required to limit host cell motility. Therefore, we added chloramphenicol at 256 - 512 μg/ml, a concentration known to prevent the synthesis of the Cia effector proteins ^32; 62^, resulting in a decrease in bacterial invasion ^32^. We observed a significant decrease in cell processivity with the *C. jejuni* wild-type strain (Figure 9B). Infection of the A549 cells with the *C. jejuni* wild-type strain with chloramphenicol and infection with the *C. jejuni* Δ*cadF* Δ*flpA* mutant partially restored processivity. In addition, infection of the A549 cells with the *C. jejuni* Δ*cadF* Δ*flpA* mutant with chloramphenicol resulted in the highest level of processivity of all infected conditions. In conjunction with the previous finding, these results suggest that the *C. jejuni* CadF and FlpA fibronectin-binding proteins and secreted effector proteins contribute to *C. jejuni* focal adhesion modification.

Lastly, we investigated the dependence of CadF and FlpA in collective cell migration. T84 cells were grown to confluency and then scratched by aspiration to create circular wound beds. Cells were infected with *C. jejuni* wild-type strain, C. *jejuni* Δ*cadF* Δ*flpA* mutant, and the *cadF flpA* complemented isolate. We observed a significantly decreased rate of scratch closure in *C. jejuni* wild-type infected cells compared to non-infected cells (Figure 9C-D). Further, we observed that the *C. jejuni* Δ*cadF* Δ*flpA* mutant had a rate of healing similar to the non-infected cells, while the complemented isolate showed the same phenotype as the wild-type condition. Collectively, these results provide evidence that the CadF and FlpA adhesins contribute to the functional and physical changes in the focal adhesion.

## DISCUSSION

The key findings of this study show that *C. jejuni* changes the structure, composition, and function of cellular focal adhesions using a combination of virulence factors. *C. jejuni* changes the focal adhesion structure by affecting paxillin footprint size, turnover rate, and nanoscale position. Since paxillin is a dynamic regulator of focal adhesion function, these alterations dramatically change host cell behavior. This was dependent on bacterial virulence factors, including the flagellum and the CadF and FlpA fibronectin-binding adhesins, and required the host cell FAK and Src kinases. The biological outcomes of such manipulations are that cell adhesion, cell motility, and epithelial sheet migration/wound healing were significantly altered by *C. jejuni* infection. The defects in cell motility are likely due to a disruption in the balanced lifecycle of focal adhesions. We propose that the purpose of taking over focal adhesions is driven by *C. jejuni* needing to utilize structural components for invasion. We have seen these changes by imaging both fixed and live cells, and, to our knowledge, this is the first time *C. jejuni* has been shown to manipulate the structure and dynamics of the focal adhesion.

An organized balance between focal adhesion assembly and disassembly allows coordinated host cell motility. Research has shown that focal adhesion proteins exist in two states: the cytoplasmic pool, which diffuses quickly throughout the cell, and the focal adhesion, which diffuses much slower ^21^. Focal adhesion proteins must assemble from the cytosolic pool to anchor the leading edge of the cell, then disassemble and detach from the trailing edge as the cell moves ^47^. It is the coordination of these processes that allow cells to have directional migration ^20^. If focal adhesion turnover is greater than focal adhesion assembly, or if assembly is greater than turnover, cells will be non-motile ^20^. We found that *C. jejuni* increases the size of the focal adhesion, suggesting that assembly may be occurring more than turnover. This is further supported by our observation that paxillin localization changes from the cytosol to the focal adhesion during infection. We also observed the focal adhesion to be thicker at the nanoscale level, again pointing to increased protein levels at the focal adhesion. Research has shown that focal adhesion size positively correlates with cell speed until an optimum is reached, at which point it negatively correlates ^63^. Based on our cell motility results, we hypothesize that during *C. jejuni* infection, the focal adhesion has increased past this optimum size, resulting in decreased processivity. Focal adhesion assembly and disassembly is a highly regulated process. In addition to the assembly and disassembly balance, individual proteins that compose focal adhesions are constantly joining and leaving the focal adhesion, with each individual component having a unique turnover rate and residency time ^21^. We investigated the turnover of paxillin at focal adhesions to further understand how assembly was increased during *C. jejuni* invasion. Interestingly, we found that the turnover of paxillin was decreased in infected cells, suggesting that the increased size of the focal adhesion could be driven by slower turnover (longer residency time) of individual proteins. In support of this model, research has found that larger focal adhesions have an increased protein residency time ^21^. We hypothesize that *C. jejuni* increases the focal adhesion footprint size by increasing the residency time of signaling proteins such as paxillin. The most common pathway for paxillin activation is through FAK and Src kinases.

The FAK-Src signaling pathway is required for *C. jejuni* invasion and paxillin phosphorylation. FAK is a key regulator of focal adhesion dynamics, being implicated in both focal adhesion maturation and turnover ^63^. Upon integrins binding to fibronectin ^21; 50; 64^, FAK becomes activated by autophosphorylation on tyrosine 397. This creates a high-affinity SH2 docking site for Src kinase ^50; 64; 65^. Src then phosphorylates other residues on FAK, and the now active kinase complex can phosphorylate many focal adhesion proteins, including paxillin ^50^. Phosphorylated paxillin also has a higher affinity for FAK binding ^47^, creating a positive feedback loop to recruit more FAK to focal adhesions. Interestingly, previous researchers have found that the phosphorylation of paxillin can influence assembly/disassembly from the focal adhesion ^47; 66; 67^. We observed that *C. jejuni* causes the phosphorylation of paxillin at tyrosine 118. Phosphorylation at this site, as well as at tyrosine 31, promotes nascent focal complex formation^10^. We also found that *C. jejuni* infection of cells has decreased paxillin turnover at focal adhesions. These data suggest that *C. jejuni* utilizes paxillin to manipulate focal adhesions in a unique way, mixing signals for assembly and disassembly to stunt focal adhesion functions (motility, healing, turnover).

In addition to regulating focal adhesion dynamics, paxillin phosphorylation can also recruit downstream signaling intermediates such as Erk2 and CrkII ^10; 50^. The phosphorylated tyrosine residues on paxillin create high affinity SH2 binding sites for the CrkII adaptor protein ^35^. This results in membrane ruffles via activation of Rac1 by CrkII association with the GEFs DOCK180 and ELMO ^47; 68^. Rac1 has been shown to be activated during *C. jejuni* invasion of epithelial cells ^41; 69; 70^, and DOCK180 is required for maximum invasion ^51^. We hypothesize that signaling through the focal adhesion leads to membrane ruffling and subsequent invasion. Another role for focal adhesion signaling through paxillin is by acting as a scaffold for Erk. Src-FAK phosphorylation of paxillin at Y118 can lead to Erk association ^50; 71^, and the MAP kinase cascade has been shown to be activated during *C. jejuni* infection ^61; 72^. Activation of host cell signaling has been shown to be dependent on *C. jejuni* invasion factors, in particular the CiaD secreted effector; therefore, we investigated what *C. jejuni* factors were driving changes in the focal adhesion.

We investigated the potential contribution of the *C. jejuni* adhesins and secreted effectors (via a mutated flagellar apparatus) and observed that both are involved in manipulating the focal adhesion. Regarding the adhesins, it is known that the CadF and FlpA adhesins promote the binding of the bacteria to the fibronectin localized on the basolateral surfaces of cells ^73; 74^ and stimulate the cell signaling pathways necessary for cell invasion ^49; 53; 73^. We also found that a functional flagellum/secretion apparatus was required for *C. jejuni* to increase the size of the focal adhesion and limit host cell motility using Δ*flgL*, Δ*flhF*, and Δ*cheB* mutants ^32; 42^. We propose a model whereby the CadF and FlpA adhesins promote host cell contact, which permits the delivery of effectors proteins and further modification of host cell signaling pathways. This proposal is consistent with the finding that the combination of adhesin mutant and the use of chloramphenicol led to the greatest individual cell motility of all infected conditions. Further research is required to define the precise role of the adhesins and secreted proteins in the modification of the focal adhesion and their impact on individual and collective cell migration.

It is common for bacterial pathogens to target the focal adhesion, due to the nature of focal adhesions in signaling between the extracellular matrix and the cytoskeleton. *Salmonella* Typhimurium is an extensively studied intestinal pathogen that invades host cells by secreting effector proteins into the cell. Several focal adhesion proteins have been implicated in *Salmonella* invasion. *Shi* et al. 2006 found that Salmonella recruits FAK, Cas, and paxillin to sites of bacterial invasion, creating focal adhesion-like structures. Interestingly, these structures form on the apical rather than basal cell surface of polarized cells and do not include beta integrins ^31^. This is in contrast to *C. jejuni*, where we observed an increase in the size of preexisting focal adhesions rather than the creation of new focal adhesion-like structures surrounding the bacterium. Furthermore, *Shi* et al. found that FAK and Cas are required for bacterial invasion but that the catalytic domains of FAK are not ^31^. In agreement with this observation, inhibition of Src kinase by PP2 did not decrease invasion ^31^. This is again in contrast to the results we observed with *C. jejuni*, where the catalytic activity of Src and FAK are required for invasion and paxillin phosphorylation. Previous research has also shown that talin and α-actinin, structural components of the focal adhesion, assemble at sites of *Salmonella* bacterial invasion ^75^. It is still unknown what effectors are responsible for the recruitment of FAK and Cas ^76^ and how the pathway connects to invasion. However, a recent study has shown that in macrophages, SPI-2 effector proteins target FAK to avoid fusion of the *Salmonell*a containing vacuole with the lysosome ^77^. In addition, *S*. Typhimurium infection of bone marrow-derived macrophages causes irregular movement and decreased directional movement. This is dependent on Sse1, an effector of SPI-2 ^78^. This result is consistent with our finding that *Salmonella* decreased the processivity of A549 epithelial cells. While macrophages utilize podosomes rather than focal adhesions for motility, many of the regulatory proteins, such as FAK and paxillin, are involved in both ^79^.

We observed by the trypsin detachment assay that *C. jejuni* increases focal adhesion attachment strength. The epithelial cells of intestinal villi are regularly shed by cell extrusion and programmed cell death at the villi tips. New cells migrating up from the crypts then replace these lost cells. An equilibrium between division and migration from the crypts and shedding at the tips keeps the number of cells constant ^80^. Pathogens take advantage of this natural process to cause disease. *Listeria monocytogenes* invade the villi at sites of extrusion, taking advantage of the briefly available E-cadherin ^81^. Other enteric pathogens block extrusion to prevent infected cells from being shed ^82^. The OspE effector, which is conserved in *Shigella flexneri*, EPEC, EHEC, *Salmonella*, and *Citrobacter rodentium*, interacts with integrin linked kinase (ILK) to block epithelial cell turnover and increase cell adhesion to the matrix ^82; 83^. In addition, this interaction decreases wound healing, blocks focal adhesion turnover, and increases the number of focal adhesions ^82; 83^. This is consistent with our observations that *C. jejuni* increases adhesion strength, decreases turnover, and blocks migration. However, OspE interaction with ILK causes a decrease in paxillin and FAK phosphorylation, suggesting that the pathway by which *C. jejuni* causes these changes varies from the OspE pathway. Pathogens often increase host focal adhesion strength to prevent cell extrusion from the villi. This manipulation can be two-fold, as altering focal adhesion dynamics can also prevent the villi from healing.

The piglet model of *C. jejuni* disease has shown that infection results in cell necrosis and villus blunting ^26^. The process of repairing damage is termed restitution ^14; 15; 16; 17; 18^. Most research has been done on wounds created by ischemia, viral infection, parasites, or chemicals ^18^. However, villus blunting and denuding as the epithelium lifts from the basement membrane is common in all of these agents ^14; 17; 18; 84^. Several sequential steps are involved in repairing villi damage. First, the villus contracts to shed damaged cells ^14; 85^. Next, viable cells depolarize and flatten, extending lamellipodia membrane protrusions into the damaged area ^14; 15; 16; 17; 18^. These cells then migrate to cover the damaged area. This process is dependent upon focal adhesions assembling on the leading edge of the cell and disassembling to detach from the trailing end, similar to single cell motility ^14^. Finally, tight junctions are restored as the villi wound closes ^14; 16^. For *in vitro* studies, T84 cells are commonly used ^15; 85; 86^. We observed that *C. jejuni* decreases the collective migration of T84 cells (wound healing) and hypothesize that this is due to focal adhesion manipulation. In support of this hypothesis, this effect has been observed in *E. coli*. The cytotoxic necrotizing factor 1 (CNF1) from *E. coli* has been shown to decrease T84 wound healing ^86^. CNF1 has also been shown to cause paxillin and FAK phosphorylation, further supporting the idea that migration is blocked due to manipulation of focal adhesion components ^86^.

We have laid the groundwork for understanding how *C. jejuni* manipulates the focal adhesion during infection by performing a mechanistic study focusing on the functions of focal adhesions. We observed that the focal adhesion size, Z position, and thickness increase in response to infection. We observed that these phenotypes were driven by well-known *C. jejuni* virulence determinants that, together, allow the changes in focal adhesion structure and signaling function. We hypothesize that these events have biological significance in prolonging *C. jejuni* infection of the human intestine. To summarize, we have identified a new cellular phenotype in *C. jejuni*-infected cells subsequent to cellular invasion. Studies are ongoing to understand how focal adhesion manipulation is directed towards early-infection signaling to allow invasion and late-infection manipulation to prolong sickness in a host.

## MATERIALS AND METHODS

### Bacterial strains and growth conditions

*Campylobacter jejuni* strain 81-176 was cultured on Mueller-Hinton agar (Hardy Diagnostics, Santa Maria, CA, United States) containing 5% citrated bovine blood (MHB agar) under microaerobic (10.5% CO_2_) conditions at 37 °C in a Napco 8000WJ incubator (Thermo Fisher, Waltham, MA, United States). The bacteria were subcultured on MHB agar every 24 to 48 hours. Before infection of cultured epithelial cells, *C. jejuni* were grown overnight on a MHB plate overlaid with MH broth (biphasic conditions). *Salmonella enterica* serovar Typhimurium (*S*. Typhimurium) SL1344 and *Staphylococcus aureus* ATCC 25923 were cultured on LB agar and LB broth as needed. Prior to use, the *Salmonella* was diluted 1:80 and grown for 2.5 hours to reach log phase, which corresponded to an OD_600_ of ∼0.8 ^87^.

### Generation and culture of *C. jejuni* deletion mutants and complemented isolates

The *C. jejuni* 81-176 Δ*flgL*, Δ*cheB*, and Δ*flhF* mutants, and the *flgL* complemented isolate, were generated as outlined elsewhere ^32^. The *C. jejuni* 81-176 Δ*cadF* Δ*flpA* mutant and complemented isolate were generated as described elsewhere ^73^. The Δ*flgL*, Δ*flhF*, and Δ*cheB* mutant were passaged on MHB agar plates supplemented with 8 μg/mL of chloramphenicol and the *flgL* complemented isolate, Δ*cadF* Δ*flpA* mutant, and *cadF flpA* complemented isolate were passaged on MHB agar plates supplemented with 250 µg/mL of hygromycin B.

### Cell lines

Epithelial INT 407 (ATCC CCL-6), epithelial lung carcinoma A549 (ATCC CCL-185), epithelial colonic carcinoma T84 (ATCC CCL-248), and rat bladder carcinoma cell line 804G were cultured in Minimal Essential Media (MEM; Gibco, Grand Island, NY, United States) supplemented with 10% fetal bovine serum (FBS; Heat Inactivated Seradigm Premium Grade Fetal Bovine Serum, VWR, Radnor, PA) and 1 mM sodium pyruvate (Corning Inc., Manassas, VA, United States) at 37°C with 5% CO_2_. The A549 and 804G cells were a kind gift of Jonathan C. R. Jones (WSU, Pullman, WA).

### Immunofluorescence microscopy

INT 407 cells were seeded onto glass coverslips in a 24 or 6 well dish approximately 24 hours before infection. Cells were seeded at 3 × 10^4^ cells/well for 24-well dishes or 1.5 × 10^5^ cells/well for 6 well dishes. *C. jejuni* was collected from biphasic conditions and adjusted to an OD_540_ of 0.1 in MEM supplemented with 1% FBS. Cells were rinsed once with 1% FBS MEM before adding *C. jejuni* and then incubated at 37 °C in 5% CO_2_ for 60 minutes unless otherwise indicated. Following incubation, cells were fixed with 4% paraformaldehyde for 5 to 10 minutes. Cells were then rinsed and permeabilized with 0.1% Triton X-100 in PBS containing 0.3% bovine serum albumin. *C. jejuni* was stained using a polyclonal rabbit anti-*C. jejuni* antibody (1:4000), paxillin was stained with a mouse anti-paxillin antibody (1:250, BD Biosciences, San Jose, CA), and phospho-paxillin was stained with a rabbit anti-phospho-paxillin antibody (Y118, 1:50, Cell Signaling Technologies). Secondary antibodies that were used for visualization included an Alexa Fluor 680 conjugated anti-rabbit antibody (1:1000, Jackson ImmunoResearch, West Grove, PA) and an Alexa Fluor 594 conjugated anti-mouse antibody (1:1000, Jackson ImmunoResearch). Where necessary, actin was stained with FITC conjugated phalloidin (1:1000, Sigma-Aldrich, St. Louis, MO). Images of cells infected with *C. jejuni* were taken with a confocal microscope (Leica Microsystems TCS SP5 and a Leica Microsystems TCS SP8 with LIGHTNING). Images were quantified with Fiji. To measure focal adhesion size, the paxillin channel was used. The background was subtracted and a threshold was set that included only focal adhesions greater than 0.25 μm^2^. Focal adhesions from all images were combined to ensure >1000 focal adhesions were analyzed per condition. Focal adhesions greater than 7 standard deviations away from the mean were removed from subsequent analysis. For quantification of paxillin localization, the *C. jejuni* channel was removed, and images were randomized and blinded. Images were scored based on paxillin localization (0 = paxillin at the focal adhesion, 2 = paxillin in the cytosol) by a trained individual who did not help in image acquisition. For phospho-paxillin, the focal adhesion location was measured as described above, then all focal adhesions were added to the region of interest (ROI) manager. The intensity (mean gray value) of the paxillin and phospho-paxillin channel at the focal adhesions was measured from the ROI manager. Then, the phospho-paxillin intensity was divided by the paxillin intensity at each focal adhesion.

### iPALM microscopy

iPALM images were captured using a similar procedure as described previously ^46^. Briefly, INT 407 cells were seeded on a gold fiducial coated coverslip and grown overnight prior to transfection with a tdEos-Paxillin expressing plasmid using Lipofectamine 3000. Approximately 18 hours after transfection, the cells were infected with *C. jejuni* that was collected from a biphasic culture and adjusted to an OD_540_ of 0.3. The cells were infected for 45 minutes prior to fixation in 0.4% paraformaldehyde and 0.1% glutaraldehyde in PBS for 5 minutes. Samples were washed with PBS and quenched with 1% NaBH_4_. Immunostaining for *C. jejuni* was performed as described above. Images were taken with a laser power intensity of 2 kW/cm^2^ and an exposure time of 50 ms. The number of frames acquired per image was 25000. Peakselector software was used to localize data, perform drift correction, and render images.

### iPALM data processing

ImageJ was used to process the image output files. Focal adhesions were selected from two-dimensional tiff outputs of iPLAM images by smoothing the image (Gaussian blur), thresholding, and analyzing particles greater than 0.5 μm^2^ in order to generate regions of interest (ROI). The ROI data was used to process the data output from iPALM imaging in R Studio. A LOESS (locally estimated scatterplot smoothing) regression was performed on the gold fiducials. All fluorophores were normalized to their nearest neighbor of the LOESS regression. The Z position of all fluorophores within a focal adhesion, as determined by each ROI, was averaged. Focal adhesions with Z positions calculated as less than 0 nm were removed from further analysis. Then fluorophores with Z positions less than 0 nm were removed. The average Z position of all focal adhesions was determined and compared between non-infected and *C. jejuni* infected cells. The thickness of the focal adhesion was determined as the inner quartile range (75^th^ percentile minus 25^th^ percentile) of all fluorophores within a focal adhesion.

### Live cell imaging to determine cell adhesion strength

INT 407 cells were seeded onto 35 mm tissue culture petri dishes at 4.5 × 10^5^ cells/dish approximately 24 hours before infection, with three dishes per condition. For some experiments, cells were seeded in a 24-well dish at 9.4 × 10^4^ cells/well and eight wells were used per condition. *C. jejuni* from a biphasic culture was adjusted to an OD_540_ of 0.375 in 1% FBS MEM. Cells were rinsed once with 1% FBS MEM before adding the *C. jejuni*. For nocodazole treated cells, 20 nM of nocodazole in 1% FBS MEM was added to cells after rinsing. For non-infected cells, 1% FBS MEM alone was added. Cells were incubated for 45 minutes at 37 °C before imaging. Cells were rinsed once with PBS, then a low concentration (0.05%) of trypsin with 684.4 μM EDTA was added to induce cell rounding. A phase-contrast microscope (Nikon Eclipse TE2000-U) was used to capture an image every 5 to 30 seconds. Images were collected for 16 minutes. At each minute, rounded cells were identified by the Trainable Weka Segmentation Fiji plugin or Ilastik trainable image segmentation software trained to identify rounded cells. The number of rounded cells each minute was divided by the total number of cells counted in the first image.

### A549 motility

To collect the extracellular matrix, 804G cells were grown to confluency and incubated for 6 days. The supernatant (conditioned medium) was collected, spun, filtered, and frozen until use. 12 well dishes were coated with the 804G conditioned media for 2 to 4 hours, then rinsed with PBS before seeding A549 cells at a density of 7.5 × 10^3^ cells/well 16-20 hours before infection. Experiments were conducted on an automated Leica DMi8 microscope with a heated stage. *C. jejuni* were collected from a biphasic culture and the OD_540_ was adjusted to 0.1 in 10% FBS MEM. For some experiments, the 10% FBS MEM was made with reduced sodium bicarbonate (1.2 mM) and 10 mM HEPES. For some experiments with the *C. jejuni* Δ*cadF* Δ*flpA* mutant and *cadF flpA* complement isolate, cells were pre-incubated with 1 mL of 0.1% FBS MEM for 30 minutes, then 0.5 mL of 30% FBS MEM was added immediately prior to imaging to bring the total concentration of FBS to 10%. For experiments with chloramphenicol (GoldBio, St. Louis, MO), the bacteria were incubated for 30 minutes prior to cell infection with the drug to reduce invasion but not bacterial viability (256μg/ml or 512 μg/ml ^32; 88^). Chloramphenicol was maintained for the duration of the infection and imaging. Cells were imaged for five hours, with one image taken every 5 minutes. Images were processed with an in-house localization software and the TrackMate Fiji plugin. All code used to process the images is available as open-source software at https://github.com/nimne/ACIT. Statistical analysis and wind rose plots were done using R. Processivity was calculated as path distance divided by total displacement every 5 minutes, and averaged for all cells. For experiments using *S*. Typhimurium and *S. aureus*, A549 cells were preincubated with bacteria in 10% FBS MEM with 26.2 mM sodium bicarbonate for 60 minutes in a 5.5% CO_2_ 37 °C incubator. Cells were then rinsed 3 times with PBS to remove excess bacteria, then imaged as described above in low sodium bicarbonate 10% FBS MEM. Cells were infected with approximately 2.4 × 10^8^ CFU of *S*. Typhimurium and *S. aureus*. Chloramphenicol was added to prevent bacterial replication during the assay (*S*. Typhimurium = 8 µg/mL and *S. aureus* = 64 µg/mL).

### Paxillin immunoprecipitation for inhibitor experiments

INT 407 cells were seeded at a density of 6 × 10^5^ cells/well in six-well tissue culture trays and incubated at 37 °C in a humidified, 5% CO_2_ incubator overnight. Cells were rinsed with MEM lacking FBS and infected with the *C. jejuni* wild-type strain and Δ*flgL* mutant at an OD_540_ of 0.3 and incubated for 45 minutes. After the incubation, the cells were rinsed three times with ice-cold PBS and lysed by the addition of ice-cold IP lysis buffer [25 mM Tris-HCl pH 7.5, 1 mM EDTA, 50 mM NaF, 150 mM NaCl, 5% glycerol, 1% Triton X-100, 1x Protease Inhibitor Cocktail (Sigma), 1 mM Na_3_VO_4_, and 1x Halt™ Phosphatase Inhibitor Cocktail (Thermo Scientific, USA)] and incubated for 20 minutes on ice. The lysate was clarified by centrifugation at 14,000 rpm for 15 minutes at 4 °C. Immunoprecipitation was performed by incubating mouse anti-Paxillin antibody (1:250, BD Biosciences, USA) with the lysate overnight at 4 °C followed by addition of Protein A/G PLUS-Agarose beads (Santa Cruz Biotechnology, Dallas, TX) and incubation for 2 hours at 4 °C. The precipitate was rinsed four times with ice-cold IP wash buffer (20 mM HEPES, 150 mM NaCl, 50 mM NaF, 1 mM Na_3_VO_4_, 0.1% Triton X-100, 10% glycerol, 1x Protease Inhibitor Cocktail, 1 x phosphatase inhibitor cocktail). Samples were analyzed by SDS-PAGE and immunoblot analysis, as outlined previously ^49^. Immunoblot detection of phosphorylated paxillin was performed using a 1:2000 dilution of a mouse anti-phospho-paxillin (Y-118) (Cell Signaling Technology, Danvers, MA) antibody. Detection of the total pool of paxillin was performed using a 1:1000 dilution of a mouse anti-paxillin antibody and a 1:4000 dilution of a peroxidase-conjugated rabbit anti-mouse IgG.

### Paxillin phosphorylation for *cadF flpA* deletion mutant experiments

INT 407 cells were seeded at 3.6 × 10^6^ cells/well in 100 cm^2^ dishes or 9.2 × 10^4^ cells/well in 24-well dishes 20-28 hours prior to infection. Cells were rinsed with MEM lacking FBS one time and then incubated with MEM lacking FBS for 3 to 4 hours. *C. jejuni* was collected from a biphasic culture, and the OD_540_ was adjusted to 0.035. The *C. jejuni* suspension was added in a volume equal to that of the volume in the well used for the serum starve. After 60 minutes of infection, cells were treated with a lysis buffer [25 mM Tris HCl, 150mM NaCl, 5% glycerol, 1% Triton X-100, 1x protease inhibitor cocktail (Thermo Scientific), and 2x phosphatase inhibitor cocktail (Thermo Scientific)] on ice for 10 minutes before collection of cell lysates by scraping. For experiments in 24-well dishes, cells were lysed with 2x Sample Buffer on ice for 5 minutes before collection. Lysate samples were analyzed by SDS-PAGE and immunoblot analysis, as outlined previously ^49^. Immunoblot detection of phosphorylated paxillin was performed using a 1:1000 dilution of a mouse anti-phospho-paxillin (Y-118) (Cell Signaling Technology, Danvers, MA) antibody. Detection of the total paxillin was performed using a 1:1000 dilution of a mouse anti-paxillin antibody. A 1:4000 dilution of a peroxidase-conjugated rabbit anti-mouse IgG was used to detect both primary antibodies.

### Binding and internalization assays

*C. jejuni* binding and internalization assays were performed with INT 407 cells, as outlined elsewhere ^88^. All assays were performed at a multiplicity of infection (MOI) ranging between 50 and 500, and repeated a minimum of 3 times to ensure reproducibility. The reported values represent the mean counts ± standard deviations derived from quadruplicate wells. To test the effect of FAK and Src inhibition on *C. jejuni* cell invasion, INT 407 cells were pre-incubated for 30 minutes in MEM containing 5 µM of TAE 226 (Selleck Chemicals, Houston, TX) or 10 μg/mL of PP2 (Sigma) in 0.5 mL of medium. Following incubation, a 0.5 mL suspension of *C. jejuni* in MEM was added to each well and the binding and internalization assays performed using standard laboratory protocols. To determine if a drug affected the viability of the INT 407 cells, the cells were rinsed twice with PBS following inhibitor treatment, stained with 0.5% trypan blue for 5 minutes, and visualized with an inverted microscope.

### Measurement of paxillin turnover

INT 407 cells were seeded at 1.5 × 10^5^ cells/well in glass-bottomed tissue culture dishes. The next day, cells were transfected with a tdEos-Paxillin expressing plasmid using Lipofectamine 3000 transfection reagents and incubated 16-20 hours. *C. jejuni* was collected from a biphasic culture, and the OD_540_ was measured. *C. jejuni* was diluted in 10% FBS MEM with 1.2 mM sodium bicarbonate and 10 mM HEPES to an OD_540_ of 0.25. Cells were incubated with *C. jejuni* or medium alone for approximately 60 minutes before imaging. Cells were imaged with a point scanning confocal microscope with a heated stage (Leica Microsystems TCS SP8X). Individual cells were selected, and one-half of the cell was photoswitched with a 405 nm laser. The cell was imaged every five seconds for 300 seconds after photoswitching. Multiple cells were imaged to ensure greater than 50 focal adhesions were examined. Images were analyzed with Fiji. At each >0.25 μm^2^ focal adhesion, the intensity of red fluorescence was measured in every frame. The slope of the red fluorescence in the non-photoswitched half of the cell was calculated by linear regression analysis.

### Collective cell migration

T84 cells were seeded at 2.25 × 10^5^ cells well in 24-well tissue culture dishes. Cells were given fresh 10% FBS MEM every day until confluency was reached (7-12 days). Cells were scratched by aspiration with a gel-loading or 0.1 – 2 µL tip. Immediately following scratching, cells were infected with *C. jejuni. C. jejuni* from a biphasic culture was adjusted to an OD_540_ of 0.3 in 1% FBS MEM and added to cells for 3 hours. After infection, the medium was replaced with 1% FBS MEM. Scratches were imaged with a phase contrast microscope 1 - 2 times daily, and the medium was replaced every day for four days to monitor wound healing. 1% FBS MEM was used to prevent cell replication into the scratches. 6 to 12 scratches were averaged for all conditions. The reported values for linear regression slopes represent the mean counts ± SEM.

### Statistical analysis

All statistical analysis was done in GraphPad Prism v6.0a or R 3.5.1. Biological replicates were defined as experiments repeated on separate days, and one of three replicates is shown for each figure unless otherwise noted.

## ACKNOWLEDGMENTS

We thank Colby Corneau for assistance with gentamicin-protection assays and paxillin phosphorylation analysis, Lauren Breymeyer, Jenna Lounsbery, and Abigail Hicks for assistance in the analysis of the microscopy images, and Satya Khuon (AIC Janelia) for technical help and advice.

## FIGURE LEGENDS

**Supplementary Figure 1**: *Salmonella enterica* serovar Typhimurium (*S*. Typhimurium), but not *Staphylococcus aureus*, decrease single cell motility. A549 cells were infected with *S. typhimurium* or *S. aureus* for 60 minutes, rinsed with PBS, and imaged for five hours with one image taken every five minutes. Cells were tracked over time, and processivity (path distance divided by path displacement) was calculated to quantify motility. Error bars represent SEM, * indicates *p* < 0.0001 (Student’s T Test). More than 500 cells imaged per condition.

**Supplementary Figure 2:** *C. jejuni* requires a motile flagella and invasive ability to limit A549 individual cell motility. A549 cells were infected with *C. jejuni* wild-type strain and the *C. jejuni* Δ*cheB*, Δ*flhF*, and Δ*flgL* flagellar mutants. The *C. jejuni* Δ*cheB* mutant, which lacks expression of *cheB* and *cheR*, has reduced motility and chemotaxis, but normal invasion and effector secretion. This mutant resulted in host cell motility equal to the wild-type strain. Mutants that are completely non-motile and non-invasive (Δ*flhF and* Δ*flgL*) showed the most restoration of motility. By one-way ANOVA with Tukey’s multiple comparison’s test: # indicates significant difference from all other conditions (*p* < 0.001), † indicates significant difference from non-infected, Δ*flhF*, and Δ*flgL* conditions (*p* < 0.001), ‡ indicates significant difference from non-infected, wild-type, and Δ*cheB* conditions (*p* < 0.001). Error bars represent SEM of > 300 cells.

**Supplementary Figure 3:** Representative images of super-resolution iPALM imaging showing that *C. jejuni* changes the nanoscale topology of the focal adhesion. **A**. Representative iPALM image of a non-infected cell. **B**. Representative iPALM image of a *C. jejuni* infected cell. **C**. Focal adhesions from the non-infected cell shown in Panel A. The Z position of fluorophores was corrected to the Z position of gold fiducials embedded into the coverslip, as described in Materials and Methods. Z position is indicated by color scale gradient. **D**. Focal adhesions from the *C. jejuni* infected cell shown in Panel B. The focal adhesions of *C. jejuni* infected cells are displaced upwards compared to non-infected cells. Focal adhesions present in Panels A and B but not panels C and D represent focal adhesions removed from analysis for having an average Z position less than 0 nm.

**Supplementary Figure 4:** Representative blot demonstrating *C. jejuni* driven paxillin phosphorylation and dependence on the CadF and FlpA adhesins and the flagellum. Whole-cell lysates from *C. jejuni* infected INT 407 cells were analyzed by SDS-PAGE and Immunoblot. Blots were probed for total paxillin and phosphorylated paxillin (Y118).

