## Supplementary figures and images for "*Campylobacter jejuni* trigger signaling through host cell focal adhesions to inhibit cell motility and impede wound repair"

### Supplemental Figure 1

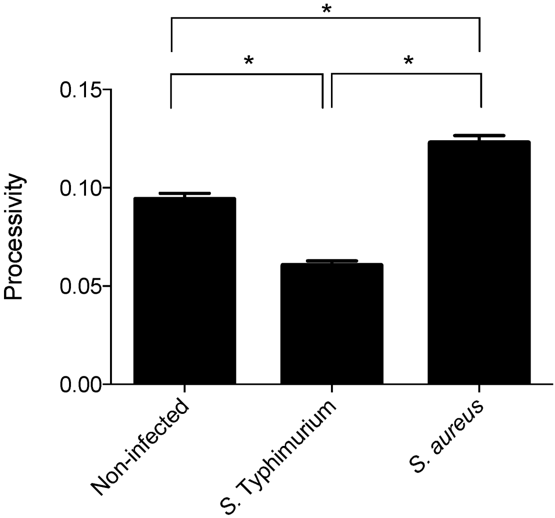

### Supplemental Figure 2

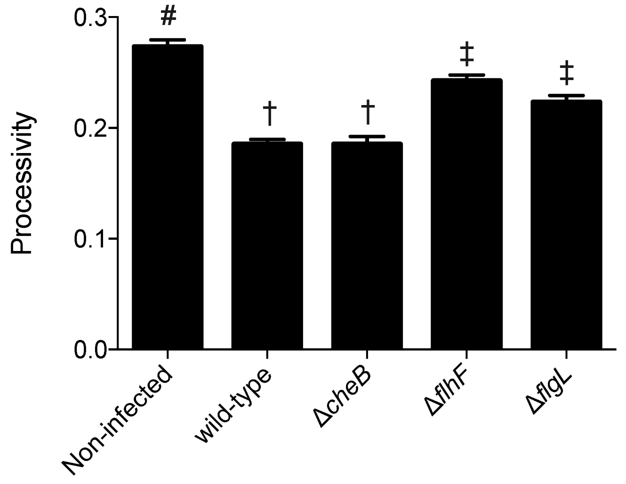

### Supplemental Figure 3

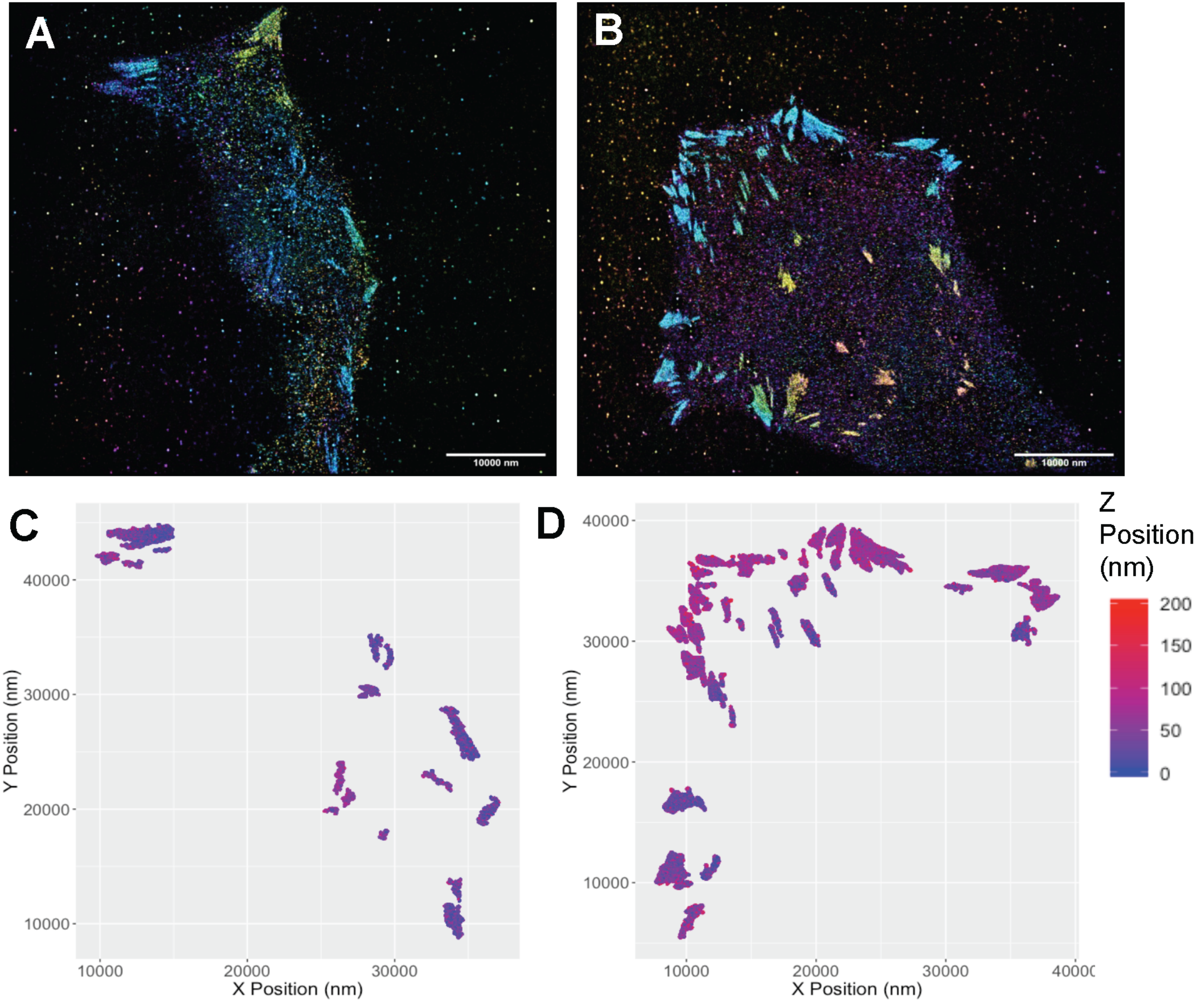

### Supplemental Figure 4

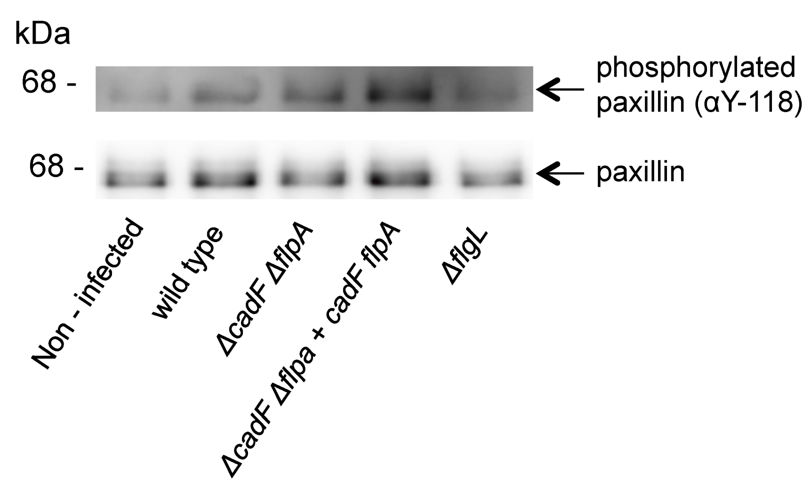
